# Asymmetrical localization of Nup107-160 subcomplex components within the nuclear pore complex in fission yeast

**DOI:** 10.1101/223131

**Authors:** Haruhiko Asakawa, Tomoko Kojidani, Hui-Ju Yang, Chizuru Ohtsuki, Hiroko Osakada, Atsushi Matsuda, Masaaki Iwamoto, Yuji Chikashige, Koji Nagao, Chikashi Obuse, Yasushi Hiraoka, Tokuko Haraguchi

## Abstract

The nuclear pore complex (NPC) forms a gateway for nucleocytoplasmic transport. The outer ring protein complex of the NPC (the Nup107-160 subcomplex in humans) is a key component for building the NPC. Nup107-160 subcomplexes are believed to be symmetrically localized on the nuclear and cytoplasmic sides of the NPC. However, in *S. pombe* immunoelectron and fluorescence microscopic analyses revealed that the homologous components of the human Nup107-160 subcomplex had an asymmetrical localization: constituent proteins spNup132 and spNup107 were present only on the nuclear side (designated the spNup132 subcomplex), while spNup131, spNup120, spNup85, spNup96, spNup37, spEly5 and spSeh1 were localized only on the cytoplasmic side (designated the spNup120 subcomplex), suggesting the complex was split into two pieces at the interface between spNup96 and spNup107. This contrasts with the symmetrical localization reported in other organisms. Fusion of spNup96 (cytoplasmic localization) with spNup107 (nuclear localization) caused cytoplasmic relocalization of spNup107. In this strain, half of the spNup132 proteins, which interact with spNup107, changed their localization to the cytoplasmic side of the NPC, leading to defects in mitotic and meiotic progression similar to an spNup132 deletion strain. These observations suggest the asymmetrical localization of the outer ring spNup132 and spNup120 subcomplexes of the NPC is necessary for normal cell cycle progression in fission yeast.

**Author summary:** The nuclear pore complexes (NPCs) form gateways to transport intracellular molecules between the nucleus and the cytoplasm across the nuclear envelope. The Nup107-160 subcomplex, that forms nuclear and cytoplasmic outer rings, is a key complex responsible for building the NPC by symmetrical localization on the nuclear and cytoplasmic sides of the nuclear pore. This structural characteristic was found in various organisms including humans and budding yeasts, and therefore believed to be common among “all” eukaryotes. However, in this paper, we revealed an asymmetrical localization of the homologous components of the human Nup107-160 subcomplex in fission yeast by immunoelectron and fluorescence microscopic analyses: in this organism, the Nup107-160 subcomplex is split into two pieces, and each of the split pieces is differentially distributed to the nuclear and cytoplasmic side of the NPC: one piece is only in the nuclear side while the other piece is only in the cytoplasmic side. This contrasts with the symmetrical localization reported in other organisms. In addition, we confirmed that the asymmetrical configuration of the outer ring structure is necessary for cell cycle progression in fission yeast. This study provides notions of diverse structures and functions of NPCs evolved in eukaryotes.

## Introduction

In eukaryotes, the nuclear envelope (NE) separates the nucleus from the cytoplasm. Molecular transport between the nucleus and cytoplasm across the NE occurs through nuclear pore complexes (NPCs). These complexes are cylindrical, eight-fold symmetrical structures that perforate the NE and are made of multiple sets of about 30 different protein species known as nucleoporins (Nups) [1–3]. Nups are classified into three groups: transmembrane Nups, FG repeat Nups, and scaffold Nups. Transmembrane Nups have transmembrane helices and anchor NPCs to the NE. FG repeat Nups contain phenylalanine-glycine (FG) rich repeats and are involved in molecular transport through the NPC cylinder structure. Scaffold Nups form two inner rings and two outer rings, which serve as the NPC structural core [4–7], and associate with the membrane through interactions with transmembrane Nups [3,8,9]. These NPC structures and most Nups are generally conserved among eukaryotes [1,2,10–13], although numerous species-dependent differences are found [14].

The Nup107-160 subcomplex is a key component of the outer rings and in most eukaryotes is composed of equal numbers of Nup107, Nup85, Nup96, Nup160, Nup133, Sec13, and Seh1, and depending on the species Nup37, Nup43, and ELYS are also included [15–20]. These Nups assemble to form, both *in vitro* and *in vivo*, the Y-shaped Nup107-160 subcomplex in *Homo sapiens*, the budding yeast *Saccharomyces cerevisiae*, and the thermophile *Chaetomium thermophilum* [5,20–25]. Nup85, Nup43, and Seh1 form one of the two short arms, while Nup160, Nup37, and ELYS form the other. The two arms are connected to Nup96 and Sec13, and Nup96 is connected to Nup107 and Nup133 to form the long stem (Nup96-Nup107-Nup133) of the Y-shaped molecule. Multiple copies of the Nup107-160 subcomplex form the outer rings on the nucleoplasmic and cytoplasmic sides of the NPC [4,5,26].

Like other eukaryotes, the fission yeast *Schizosaccharomyces pombe* has a set of conserved Nups [27–29] (hereafter, we use ‘sp’ to denote *S. pombe* proteins and ‘sc’ and ‘hs’ to indicate *S. cerevisiae* and *H. sapiens* proteins) (**S1 Table**). However, the *S. pombe* NPC has several unique features. It contains spEly5 (a potential homolog of metazoan ELYS) and spNup37, but lacks Sec13 and Nup43 [29], and it has two redundant Nup133 homologs (spNup131 and spNup132) in addition to two redundant scNic96/hsNup93 homologs (spNpp106 and spNup97). Most strikingly, the Nup107-160 subcomplex in *S. pombe* is composed of unequal numbers of spNup107, spNup120 (a homolog of hsNup160), spNup85, spNup96, spNup37, spEly5, spSeh1, spNup131 and spNup132 [29]; the relative numbers of spNup107and spNup131 are 4 - 8 in a single NPC, and that of spNup132 is approximately 48 whereas the other components are approximately16. Thus, a unique/different structural organization of the Nup107-160 subcomplex is suggested in *S. pombe* compared with *S. cerevisiae* and humans [29].

spNup131 (spNup133a) and spNup132 (spNup133b) have similar molecular structure and both are able to bind to spNup107 [27]. Despite their similar biochemical features, spNup131 and spNup132 are likely to have different functions because gene disruption strains show different phenotypes. The strain lacking spNup132 (*nup132*Δ) displays altered NPC distribution [27]; its growth is inhibited in the presence of thiabendazole (a microtubule-depolymerizing drug) or hydroxyurea (a DNA replication inhibitor) [28,29]; and it exhibits delayed chromosome segregation in meiosis and unusual spore formation [29,30]. The strain lacking spNup131 (*nup131*Δ) doesn’t display any of these characteristics. In addition, telomere elongation and deficiency in SUMOylation have been reported in *nup132*Δ-specific phenotypes [31,32]. The causes of these functional differences remain unknown.

In the present study, we examined the positioning of each Nup within the NPC in *S. pombe* using immunoelectron microscopy and high-precision distance measurements using fluorescence microscopy and found asymmetrical positionings of the outer ring complex components in the nuclear and cytoplasmic sides of the NPC. In addition, genetical alteration of positioning of the key molecule Nup132 of the outer ring subcomplex resulted in the defects observed in *nup132*Δ.

## Results

### spNup131 and spNup132 are differently positioned at the cytoplasmic and nucleoplasmic sides of the NPC

The molecular architectures of spNup131 and spNup132 are similar to that found in other species, with an N-terminal β-propeller domain followed by a C-terminal α-helix stack domain [33] (**S1A Fig**). However, phylogenetic analysis indicates that spNup131 and spNup132 belong to evolutionarily distant clades, with the spNup131-containing clade branching from a common ancestor of yeast Nup133 homologs much earlier than that of spNup132 (**S1B Fig**). This suggests that despite the similarity in their domain architecture, spNup131 and spNup132 may have structural features that are distinct enough to confer different functions.

To understand the differences between spNup131 and spNup132, we examined the positioning of those Nups within the NPC by immunoelectron microscopy (IEM). The spNup131 or spNup132 gene was replaced with the respective gene N-terminally fused to GFP (GFP-spNup131 or GFP-spNup132). IEM was carried out using a specific antibody against GFP. The results showed that GFP-spNup131 is located at the cytoplasmic side of the NPC, while GFP-spNup132 is located at the nuclear side (**Fig 1A**). To confirm the accessibility of the nucleus to immunogold particles using this method, the nuclear centromere protein spMis6 [34] was co-stained (**S2A-C Fig**); only the cells positive for spMis6 were evaluated for staining of spNup131(**S2B,C Fig**). A montage picture of spNup131 with quantification shows that spNup131 was exclusively located in the cytoplasmic side of the NPC (left panels of **Fig 1B**). In contrast, the localization of spNup132 was exclusively in the nuclear side (right panels of **Fig 1B**), indicating that spNup131 and spNup132 have distinct localizations. To resolve potential artifacts of GFP-tagging at the N-terminus, we repeated these experiments using strains in which spNup131 and spNup132 were C-terminally fused to GFP. We obtained essentially the same results with N- and C-tagged proteins (**Fig 1C**), suggesting that spNup131 and spNup132 are differentially positioned at the cytoplasmic and nuclear sides, respectively, of the NPC.

**Fig 1.**
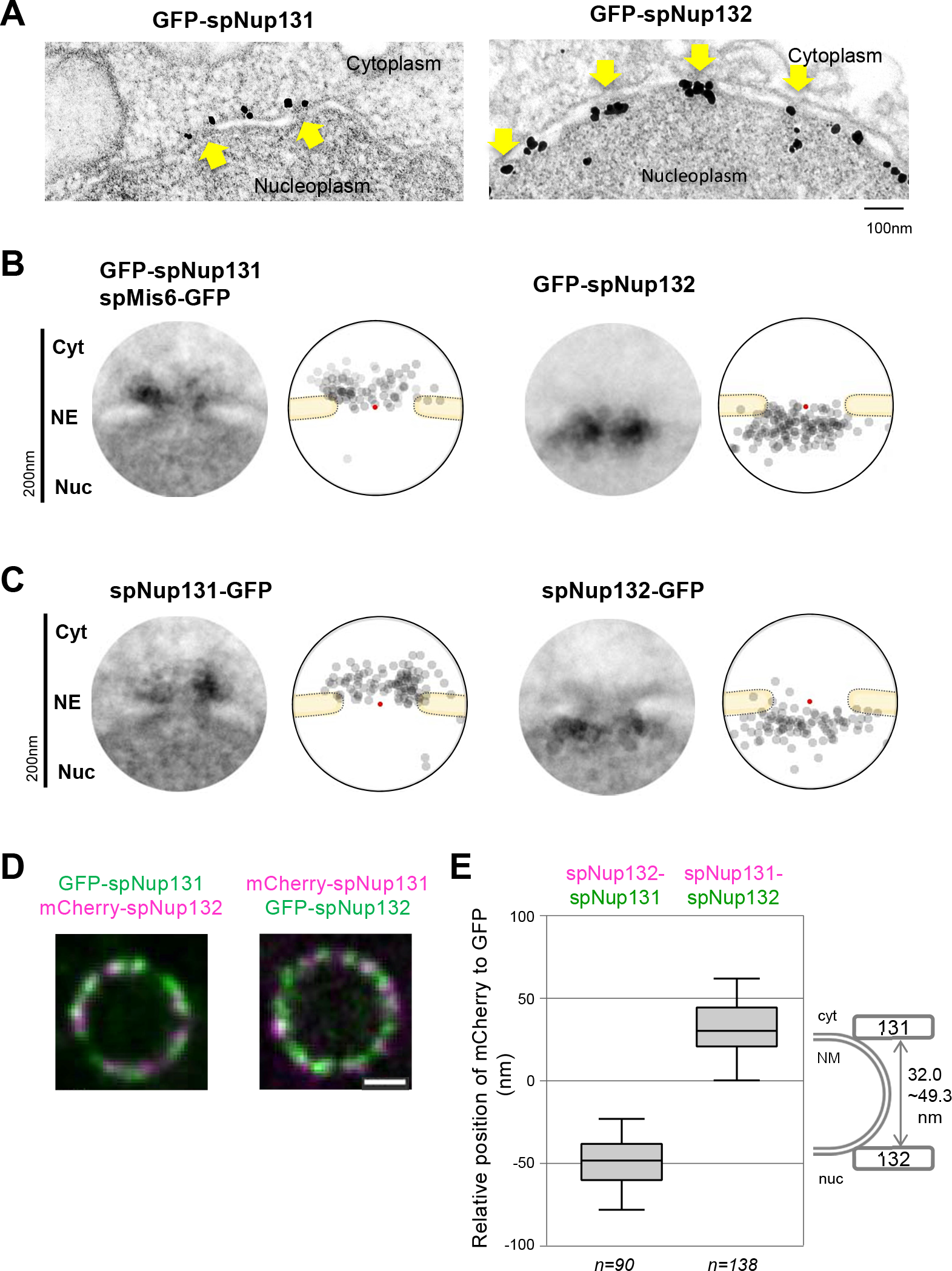
Localization of spNup131 and spNup132 at the NPC. **(A)** IEM of GFP-spNup131 and GFP-spNup132. Arrows indicate the nuclear pores. Scale bar, 100 nm. **(B)** Quantitative representation of IEM for N-terminally tagged spNup131 and spNup132. (left) A montage image of 20 immunoelectron micrographs. The diameter of the circle is 200 nm. (right) A schematic drawing illustrating the distribution of immunogold particles shown in the montage image. Red dots represent the pore centers. **(C)** Quantitative representation of IEM for C-terminally tagged spNup131 and spNup132. Montage pictures and distributions of immunogold particles are shown as described in (**B)**. IEM micrographs used for the quantitative analysis are available in S1 Dataset. **(D)** Fluorescence microscopy of a nucleus simultaneously expressing spNup131 and spNup132 fused to GFP and mCherry in living cells. Single section images of the same focal plane are shown. (left) A nucleus expressing GFP-spNup131 and mCherry-spNup132. Green, GFP-spNup131; Magenta, mCherry-spNup132. (right) A nucleus expressing mCherry-spNup131 and GFP-spNup132. Magenta, mCherry-spNup132; Green, GFP-spNup131. Scale bar, 1 μm. **(E)** Distances between spNup131 and spNup132 in living cells. Results from cells expressing GFP-spNup131 and mCherry-spNup132 (n=90, left) and results from cells expressing mCherry-spNup131 and GFP-spNup132 (n=138, right) are shown in the box plot: values of the distance determined in individual cells are shown in S2 Dataset. Center lines show the medians; box limits indicate the 25th and 75th percentiles. Whiskers extend to the 5th and 95th percentiles. The diagram on the right shows the positions of spNup131 and spNup132 within the NPC.

To confirm the different localization of spNup131 and spNup132 in living cells by fluorescence microscopy (FM), we observed cells simultaneously expressing spNup131 and spNup132 fused with GFP and mCherry, respectively; we also tested cells expressing spNup131 and spNup132 fused with mCherry and GFP, respectively (**Fig 1D**). To determine their localizations within nuclear pores with high-precision, we applied an open-source program “Chromagnon” [35] (This software is available at https://github.com/macronucleus/Chromagnon [36]). After the chromatic correction, the average distance of spNup131 from spNup132 in each nucleus along its radial direction was measured (see Materials and Methods for details). As a result, the position of mCherry-spNup132 relative to GFP-spNup131 was −49.3 ± 1.8 nm (mean ± SEM), and the position of mCherry-spNup131 relative to GFP-spNup132 was 32.7 ± 1.4 nm, both indicating that spNup131 was located at the exterior position (distant from the nuclear center) compared with the location of spNup132 within the NPC (**Fig 1E**). These results support the interpretation of the IEM observation that spNup131and spNup132 are separately localized in the NPC: spNup131 is positioned at the cytoplasmic side and spNup132 is positioned at the nucleoplasmic side of the NPC.

### Different functions of spNup131 and spNup132 at the cytoplasmic and nucleoplasmic sides of the NPC

To determine the function of spNup131 and spNup132, we identified its interacting proteins using affinity capture/mass spectrometry (**S3A, B Fig)**. Several non-Nup proteins that interact with spNup131 and spNup132 were identified (**S3C Fig**).

Among the candidate proteins specifically interacting with spNup131, we selected spFar8 (also known as spCsc3) and examined its functional relationship with spNup131. spFar8 is one of the components of the striatin-interacting phosphatase and kinase (STRIPAK) complex [37,38] that regulates the functions of the spindle pole body (SPB; the yeast microtubule-organizing center) during mitosis [39]. GFP-fused spFar8 (spFar8-GFP) localized at the nuclear periphery during interphase, as previously reported [39] (see “wild type” in **Fig 2A**). IEM analysis revealed that spFar8-GFP localized at the cytoplasmic side of the nuclear pores (**Fig 2B**). This NPC localization of spFar8-GFP was greatly decreased in the *nup131*Δ cells (see “*nup131*Δ” in **Fig 2A**) but not in the *nup132*Δ cells (“*nup132*Δ” in **Fig 2A**). Previous studies have reported that in the *nup132*Δ background, NPCs cluster on the NE [27]. This NPC-clustering phenotype in the *S. pombe nup132*Δ cells is of low penetrance, when the cells are in exponential growth. In our experiment, NPCs did not cluster in the *nup132*Δ cells because the cells were growing exponentially. Western blotting analysis of the cell strains tested above showed no marked changes in the amount of spFar8 protein (**Fig 2C**). The mislocalization of spFar8 in the *nup131*Δ cells occurred when spNup132 was ectopically expressed, whereas normal localization was restored when spNup131 was expressed (**Fig 2D, E**). Localization of another STRIPAK complex protein, spFar11, to the nuclear periphery was also decreased in *nup131*Δ cells (**S4 Fig**). These results suggest that spNup131 plays a role in retaining the STRIPAK complex at the cytoplasmic side of the NPC in interphase cells. The genetic interaction of STRIPAK proteins with spNup131 shown by this experiment is consistent with their localization at the cytoplasmic side of the NPC indicated by IEM.

**Fig 2.**
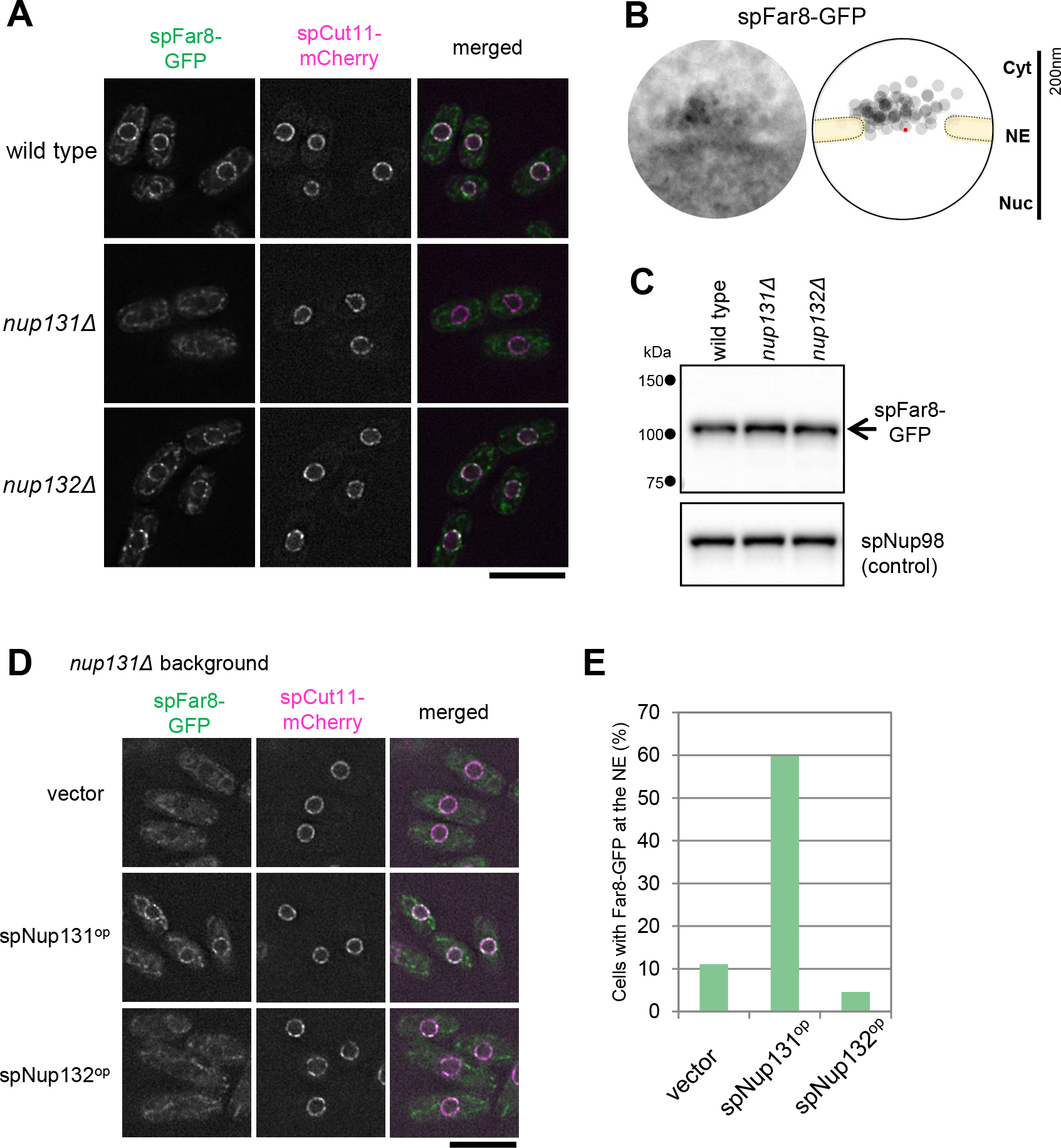
spNup131 is required for spFar8 localization on the cytoplasmic side of the NPC. **(A)** Localization of spFar8-GFP in wild type, *nup131*Δ, and *nup132*Δ cells. spFar8-GFP was expressed in the indicated strains, and cells exponentially growing in EMM2 liquid medium were observed by FM. Single section images of the same focal plane are shown. spCut11-mCherry was observed as an NPC marker. Scale bar, 10 μm. **(B)** IEM of spFar8-GFP. For quantitative representation, a montage image of 20 immunoelectron micrographs and a schematic drawing illustrating the distribution of immunogold particles are shown. The red point indicates the pore center. IEM micrographs used for the quantitative analysis are available in S3 Dataset. **(C)** Western blot analysis of spFar8-GFP. Whole cell extracts were prepared from wild type, *nup131*Δ, and *nup132*Δ cells expressing spFar8-GFP and subjected to SDS-PAGE and Western blot analysis. spFar8-GFP was detected with anti-GFP antibody. spNup98 was detected with anti-Nup98 antibody (13C2) as a loading control. The arrow indicates the position of spFar8-GFP. Small dots on the left indicate positions of molecular weight markers. **(D)** Overexpression of spNup131 and spNup132 in *nup131*Δ spNup131 or spNup132 were overexpressed in *nup131*Δ cells expressing spFar8-GFP. cells and localization of spFar8-GFP and spCut11-mCherry was observed by FM. Empty vector was also introduced for a control strain. Single section images of the same focal plane are shown. Scale bar, 10μm. **(E)** Quantitative analysis of cells exhibiting spFar8-GFP at the NE in the experiment described in **(D)**. Cells observed were: 81, 102, and 130 for vector, spNup131^op^, and spNup132^op^, respectively.

Among the candidate spNup132 interacting proteins identified by affinity capture/mass spectrometry (**S3C Fig**), we examined the functional relationship of spNup211 with spNup132. spNup211 is an *S. pombe* homolog of human Tpr and *S. cerevisiae* Mlp1 and Mlp2 [40]; Tpr homologs are known to localize on the nuclear side of the NPC [41,42]. IEM analysis of spNup211, whose C-terminal was tagged with GFP (spNup211-GFP), showed that spNup211 localized at the nuclear side of the nuclear pores as expected (**Fig 3A**). To examine its localization in relationship to spNup132, spNup211-GFP was observed by FM in wild type, *nup131*Δ, and *nup132*Δ cells. In wild type and *nup131*Δ cells, spNup211-GFP was localized at the nuclear periphery (see “wild type” and “*nup131*Δ” in **Fig 3B**). In contrast, in the *nup132*Δ cells, spNup211-GFP formed several bright foci at the nuclear periphery (“*nup132*Δ” in **Fig 3B**). We measured the maximum fluorescence intensity of spNup211-GFP in each nucleus in wild type, *nup131*Δ, and *nup132*Δ cells. The value was significantly higher in the *nup132*Δ cells (**Fig 3C, left graph**); the value was decreased when spNup132 was expressed (**Fig 3C, left graph**). On the other hand, spCut11-mCherry, as a control, was uniformly distributed at the nuclear periphery with similar maximum fluorescence intensities in all strains (**Fig 3C, right graph**). This result suggests that spNup132, but not spNup131, is the NPC component that contributes to spNup211 localization; however, we cannot exclude the contribution of additional factors. This fact is consistent with their localizations at the nucleoplasmic side of the NPC, and the lack of interaction between spNup211 and spNup131 is also consistent with their different localizations within the NPC.

**Fig 3.**
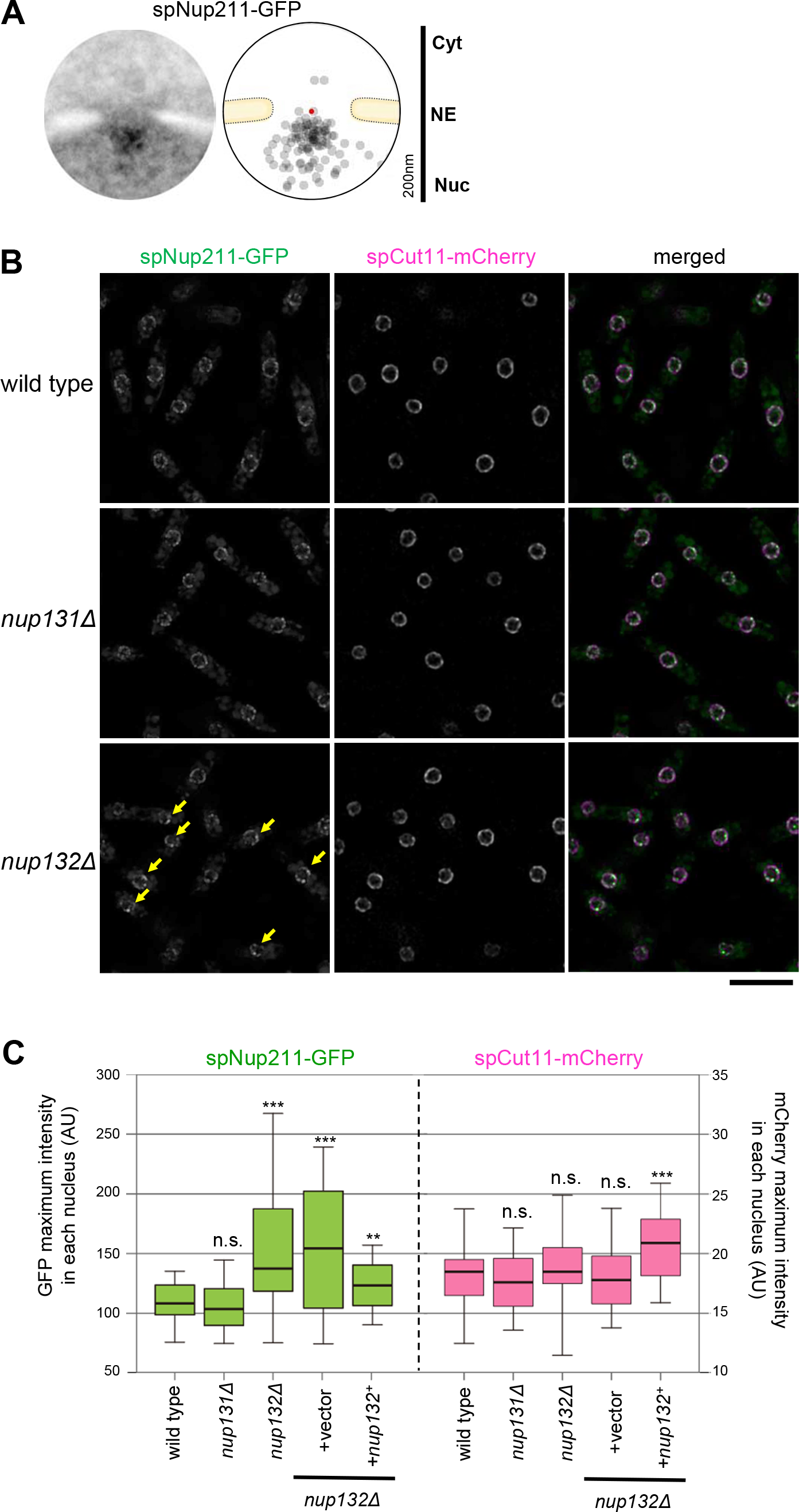
spNup132 is required for the distribution of the nucleoplasmic nucleoporin spNup211 at the NPC. **(A)** IEM of spNup211-GFP. For quantitative representation, a schematic drawing illustrating the distribution of immunogold particles based on 20 immunoelectron micrographs is shown. The red point indicates the pore center. Individual IEM micrographs and the projection image are available in S4 Dataset. **(B)** FM of spNup211-GFP in living cells growing exponentially. Single section images of the same focal plane are shown. Arrows indicate bright dots observed in *nup132*Δ cells. Scale bar, 10 μm. **(C)** Maximum intensity values of spNup211-GFP and spCut11-mCherry in wild type (n=76), *nup131*Δ (n=50), *nup132*Δ (n=73), *nup132*Δ +vector (n=61) and *nup132*Δ +*nup132*^+^ (n=46) cells: values of the maximum intensity determined in individual cells are shown in S5 Dataset. In the box plot, the center lines show the medians; box limits indicate the 25th and 75th percentiles. Whiskers extend to the 5th and 95th percentiles. Averages (± standard deviations) are 110.1 ± 22.7, 105.0 ± 21.5, 154.7 ± 64.1, 153.2 ± 56.5 and 121.9 ± 23.9 for spNup211-GFP, and 18.0 ± 3.2, 17.6 ± 3.0, 18.8 ± 4.0, 18.2 ± 3.1 and 20.8 ± 3.3 for spCut11-mCherry in wild type, *nup131*Δ, *nup132*Δ, +vector and *nup132* +*nup132*^+^, respectively. Asterisks represent the *p* value of *nup132*Δ Δ Mann-Whitney’s U-test, when compared with wild type: **, *p*<0.01; ***, *p*<0.001. n.s., no significant difference (*p*>0.05).

### Immunoelectron microscopy of spNup107-160 subcomplex Nups

Because the Nup133 homologs are integrated components of the Nup107-160 subcomplex, we next examined the positioning of the other Nup107-160 subcomplex components in *S. pombe*: spNup107 (scNup84/hsNup107), spNup120 (scNup120/hsNup160), spNup85 (scNup85/hsNup85), spNup96 (also known as spNup189C; scNup145C/hsNup96), spNup37 (hsNup37), spEly5 (hsELYS), and spSeh1 (scSeh1/hsSeh1) in cells expressing each of these Nups fused to GFP. A spNup98-spNup96 fusion protein is expressed as the *nup189*^+^ gene product, and spNup96 is generated by cleavage with the autopeptidase activity in the C-terminus of spNup98 [43].

IEM results showed that of the 7 Nups, spNup120, spNup85, spNup96, spNup37, spEly5, and spSeh1 were predominantly located on the cytoplasmic side of the NPC, whereas spNup107 was located on the nuclear side of the NPC (**Fig 4A**). The localization of spNup107 on the nuclear side was also confirmed using an N-terminus fusion protein (**Fig 4A**). Simultaneous detection of spMis6-GFP further confirmed the localization of spNup120, spNup85, spNup96, spNup37, spEly5 and spSeh1 on the cytoplasmic side of the NPC (**Fig 4A**). This result suggests that the Nup107-160 subcomplex is split into two pieces in *S. pombe* and that these two pieces are differentially located on the cytoplasmic and nuclear sides of the NPC. This result contrasts with the structure of the Nup107-160 complex reported in *S. cerevisiae* and humans, in which all of the components are connected to form the Y-shape structure. Thus, this result suggests an unexpected separation of the Nup107-160 subcomplex at the junction of spNup96 and spNup107 in *S. pombe*.

**Fig 4.**
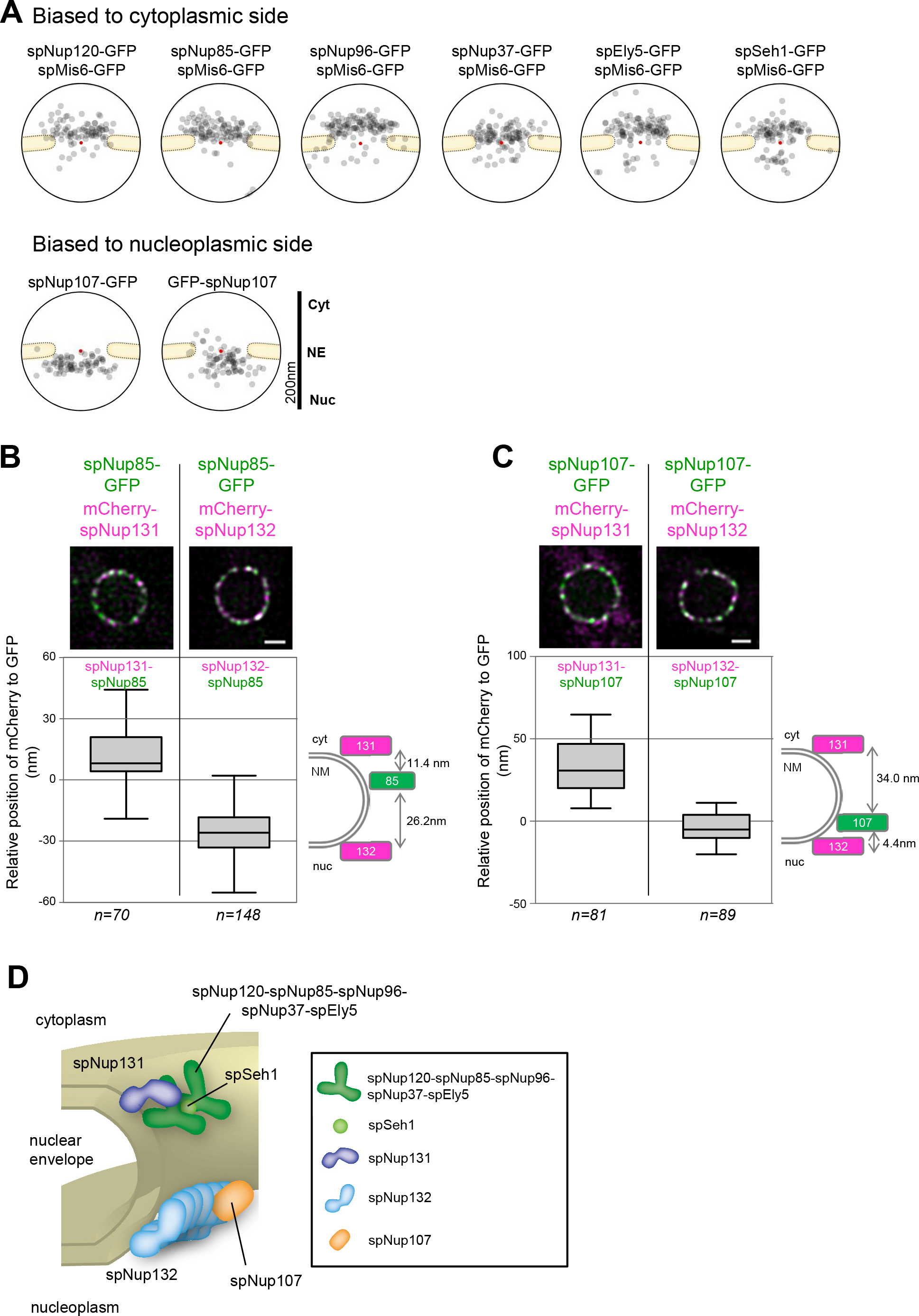
Localization of Nup107-160 subcomplex orthologs. **(A)** IEM of *S. pombe* Nup107-160 subcomplex orthologs. Immunogold distributions of the projected immunoelectron micrographs are shown as a quantitative representation. To confirm the accessibility of the nucleus to immunogold particles, spMis6-GFP was used. spMis6-positive nuclei were evaluated for staining with anti-GFP antibody. Projections of raw IEM micrographs are shown in S5A Fig. Individual IEM micrographs of 20 NPCs are available in S6 Dataset. **(B)** Position of spNup85-GFP determined by high-precision distance measurements using FM. **(upper panel)** FM images of a nucleus expressing spNup85-GFP and mCherry-spNup131 or mCherry–spNup132. Green, spNup85-GFP; magenta, mCherry-spNup131 or mCherry-spNup132. Single section images of the same focal plane are shown. Scale bar, 1 μm. **(lower panel)** Distance between spNup85-GFP and mCherry-spNup131 (left) or mCherry-spNup132 (right). The relative positions of spNup131 and spNup132 to spNup85 are shown in the box plot: center lines show the medians; box limits indicate the 25th and 75th percentiles; whiskers extend to the 5th and 95th percentiles. n, cell number examined. Values of the distance determined in individual cells are shown in S7 Dataset. The diagram on the right shows the positions of spNup85, spNup131 and spNup132 within the NPC. **(C)** Position of spNup107-GFP determined by high-precision distance measurements using FM. **(upper panel)** FM images of a nucleus expressing spNup107-GFP and mCherry-spNup131 or mCherry–spNup132. Green, spNup107-GFP; magenta, mCherry-spNup131 or mCherry-spNup132. Single section images of the same focal plane are shown. Scale bar, 1 μm. **(lower panel)** Distance between spNup107-GFP and mCherry-spNup131 (left) or mCherry-spNup132 (right). The relative positions of spNup131 and spNup132 to spNup107 are shown in the box plot as described in **(B)**. Values of the distance determined in individual cells are shown in S8 Dataset. The diagram on the right shows the positions of spNup107, spNup131 and spNup132 within the NPC. **(D)** A schematic drawing showing the localization of Nup107-160 subcomplex orthologs in *S. pombe*.

To confirm the different localizations of the Nup107-160 subcomplex components by FM, we examined the localization of these components in living cells. We chose spNup85 and spNup107 as representatives of cytoplasmic and nucleoplasmic components identified by IEM. We first determined the position of spNup85-GFP within NPCs by comparing it with that of mCherry-spNup131 and mCherry-spNup132 (**Fig 4B**). The position of mCherry-spNup131 relative to spNup85-GFP was 11.4 ± 1.9 nm (mean ± SEM) and the position of mCherry-spNup132 relative to spNup85-GFP was −26.2 ± 1.0 nm (**Fig 4B**; the relative position of Nup85 to Nup131 and Nup132 is indicated in the right panel). This result supports cytoplasmic positioning of spNup85 as indicated by IEM. Second, we determined the position of spNup107-GFP within NPCs using a similar method. The position of mCherry-spNup131 relative to spNup107-GFP was 34.0 ± 2.1 nm and the position of mCherry-spNup132 relative to spNup107-GFP was −4.4 ± 1.0 nm (**Fig 4C**; the relative position of Nup107 to Nup131 and Nup132 is indicated in the right panel). This result supports nuclear positioning of spNup107 as indicated by IEM.

Taken together, these results demonstrate that the Nup107-160 subcomplex components are differentially localized at the cytoplasmic and nucleoplasmic sides within the NPC in *S. pombe* (**Fig 4D**). Thus, we call hereafter the cytoplasmic components (spNup131, spNup120, spNup85, spNup96, spNup37, spEly5 and spSeh1) as the spNup120 subcomplex and the nucleoplasmic components (spNup132 and spNup107) as the spNup132 subcomplex.

Based on the localization analysis in this study and quantification of each Nup reported previously [29], we estimated the molecular weight of the cytoplasmic outer ring (composed of the spNup120 subcomplex) and nucleoplasmic outer ring (composed of the spNup132 subcomplex) as 7.4 MDa and 7.0 MDa, respectively: the cytoplasmic outer ring is estimated as 8-fold of a single unit (two each of spNup120, spNup85, spNup96, spEly5, and spNup37 and one each of spSeh1 and spNup131), and the nucleoplasmic outer ring is estimated as 8-fold of a single unit (six spNup132 and one spNup107). In this estimation, a single unit of both the cytoplasmic and nucleoplasmic outer rings consist of 7 α-solenoids and 6 β-propellers as elements, suggesting that similar ring structures may be consequently built in both sides. The values are a bit larger than the molecular weight (4.65 MDa) of one outer ring in *S. cerevisiae*, which consists of 8-fold of a single unit containing 5 α-solenoids and 4 β-propellers as elements [26].

### Enforced cytoplasmic localization of Nup107 by fusing to Nup96 results in reduction of Nup132 from the nuclear side, leading to defects in meiosis

To address the significance of the separated localization of the Nup107-160 subcomplex, we generated an *S. pombe* strain with spNup96 (one of the cytoplasmic outer ring Nup120 subcomplex components) artificially fused to spNup107 (one of the nuclear outer ring Nup132 subcomplex components) in a *nup107*Δ background (spNup96-spNup107-GFP). The strain was viable, and Western blot analysis confirmed expression of the fusion protein with the predicted molecular weight (**Fig 5A**). By IEM, the majority of the spNup96-spNup107-GFP fusion protein molecules were localized at the cytoplasmic side of the NPC (**Fig 5B**), showing a change in the location of spNup107 from the nuclear side to the cytoplasmic side. Under this condition, spNup132 was recruited to both the nuclear and cytoplasmic sides of the NPC (**Fig 5C**), suggesting that a significant fraction of the spNup132 molecules was recruited by spNup107. We assumed that the cytoplasmic recruitment of spNup132 molecules would result in a reduction of spNup132 at the nuclear side. It is known that the *nup132*Δ cells exhibit growth sensitivity to the microtubule-destabilizing drug, thiabendazole (TBZ) [28, 29], and also exhibit delayed meiotic division and abnormal spore formation [30]; however, the *nup131*Δ cells did not exhibit these phenotypes [29] (S6 Fig). Thus, we examined whether the spNup96-spNup107-GFP strain showed those deficiencies. We found that this strain exhibited growth sensitivity to TBZ (**Fig 5D)**, delayed meiotic division (**Fig 5E, F**), and abnormal spore formation (**Fig 5G**), similarly to the defects found in *nup132*Δ cells. This result suggests that these defects are caused by reduction in the number of spNup132 located at the nuclear outer ring of the NPC, and also suggests that concomitant localization of spNup107 and a part of Nup132 with the spNup120 subcomplex at the cytoplasmic side of the NPC caused defects in normal progression of mitosis and meiosis. Alternatively, these defects might be caused by the altered dynamics of spNup96 and spNup107 at the NPC. Thus, split structure and asymmetrical localization of the Nup107-160 subcomplex are necessary for the normal progression of mitosis and meiosis in *S. pombe*.

**Fig 5.**
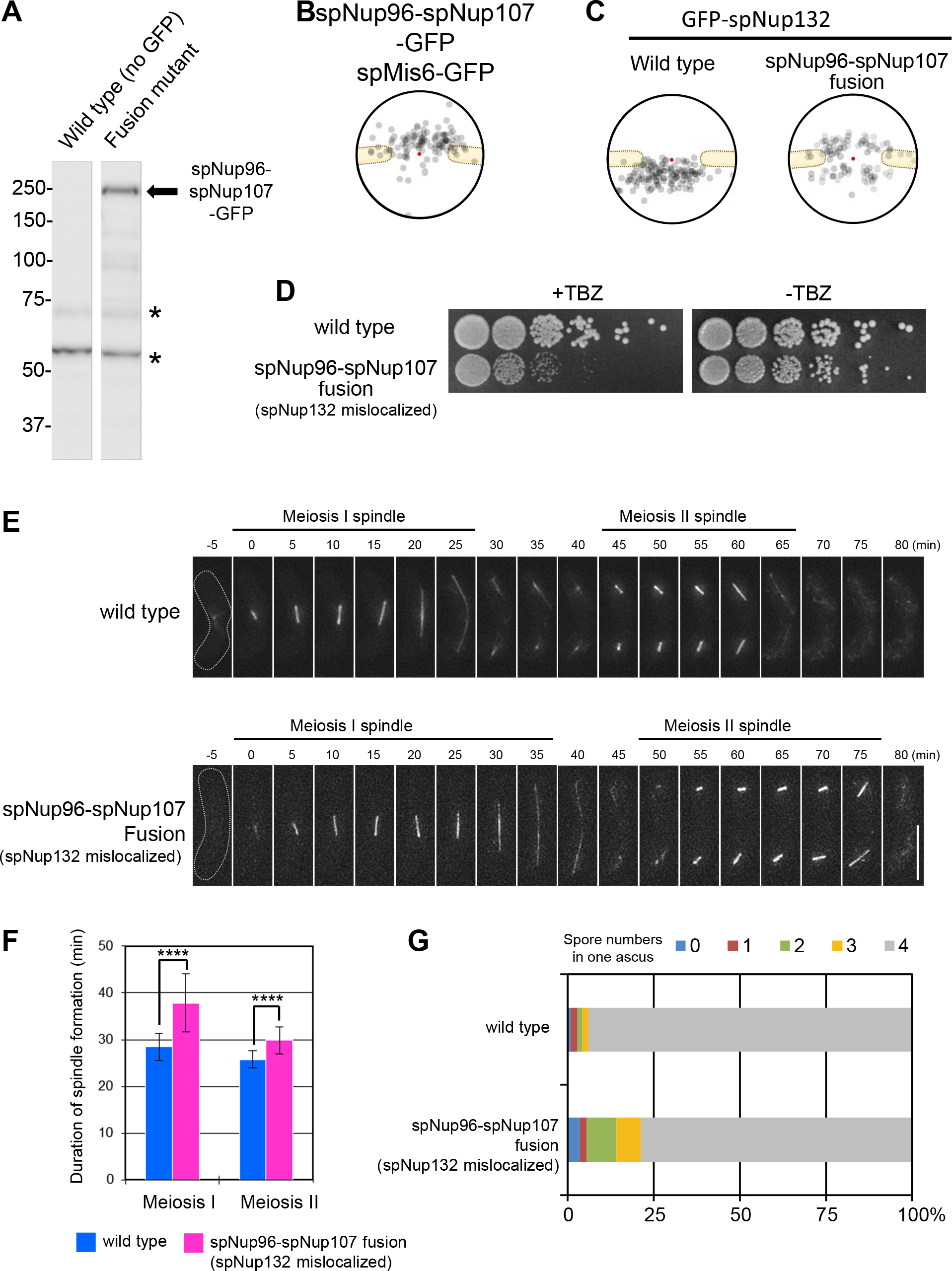
Localization and functional analysis of an spNup96-spNup107 fusion Nup. **(A)** Western blot analysis of the spNup96-spNup107-GFP fusion protein. Asterisks represent non-specific cross reactions of the anti-GFP antibody. **(B)** IEM of the spNup96-spNup107 fusion protein. Immunogold distribution of the projected immunoelectron micrographs is shown. Projection of raw IEM images is shown in S5B Fig. Individual IEM micrographs of 20 NPCs are available in S9 Dataset. **(C)** IEM of GFP-spNup132 in wild type and spNup96-spNup107 fusion backgrounds. Localization of GFP-spNup132 in a wild type background is taken from the data shown in **Fig 1B**. Projection of raw IEM images is shown in S5B Fig. Individual IEM micrographs of 20 NPCs are available in S9 Dataset. **(D)** A cell growth assay in the presence (+TBZ) or absence (−TBZ) of TBZ. Five-fold serial dilutions of wild type and spNup96-spNup107 fusion strains were spotted on YES medium containing or lacking TBZ and incubated for 3 days. **(E)** Time-lapse observation of *S. pombe* cells (wt or spNup96-spNup107 fusion strains) undergoing meiosis. Cells expressing mCherry-fused α-tubulin (spAtb2) were induced to enter meiosis. The duration of meiosis I and II was determined as the time the spindle was present. Dotted lines show cell shapes. The time 0 indicates the time of the first appearance of meiosis I spindle formation. Representative images are shown as a maximum intensity projection of the serial z-sections (number of cells observed: 32 for wild type and 33 for spNup96-spNup107). **(F)** Statistical analysis for images obtained in **(E)**. Duration of meiosis I and II. Error bars represent standard deviations. The duration of meiosis I was 28.4 ± 3.0 min in wild type and 37.9 ± 6.3 min in spNup96-spNup107 fusion cells. The duration of meiosis II was 25.8 ± 1.8 min in wild type and 29.8 ± 2.9 min in spNup96-spNup107 fusion cells. Asterisks indicate statistical significance (*p* < 0.0001) between the indicated strains by Welch’s t-test. Number of cells observed: 32 for wild type and 33 for spNup96-spNup107. **(G)** Abnormal spore formation was observed in the spNup96-spNup107 fusion background. More than 200 asci were counted for each strain.

### The N-terminal β-propeller region is required for differential localization of Nup133 homologs

Finally, we determined which domains of spNup131 and spNup132 were responsible for their different localizations. In this experiment, we expressed full length spNup131 or spNup132, fragments of spNup131 or spNup132, or chimeric proteins of spNup131 and spNup132 (**Fig 6A**). To exclude the possibility that endogenous spNup131 or spNup132 predominantly occupy preferred sites for spNup131 and spNup132 fragments or chimeric proteins, a *nup131*Δ *nup132*Δ double-deletion strain (lacking genes for both spNup131 and spNup132) was used. Protein expression was confirmed by Western blot analysis (**S7A Fig**). FM observation showed that GFP fused full-length spNup131 (spNup131FL) and spNup132 (spNup132FL) were localized at the nuclear periphery (**Fig 6B**). The N-terminal domains of both Nup131 and Nup132 (spNup131N and spNup132N) were not localized at the nuclear periphery, while the C-terminal domain of both proteins (spNup131C and spNup132C) were localized at the nuclear periphery, consistent with a previous report [44]. Interestingly, IEM analysis showed that spNup131C localized to the nuclear side of the NPC (**Fig 6C**) in contrast with the localization of full length spNup131 on the cytoplasmic side (compare the upper two panels in **Fig 6C**; also see **Fig 1B**). These results suggest that the C-terminal domains of both spNup131 and spNup132 have the ability to localize on the nuclear side of the NPC, probably through binding to Nup107, and also suggest that the N-terminal domains of spNup131 and spNup132 are involved in determining their differential localizations. To test this idea, we examined the localization of the chimeric proteins spNup131N+spNup132C and spNup132N+spNup131C: If the C-terminal domains were independent and functioning correctly, both chimeric proteins should be localized to the nuclear side of the NPC via interaction with Nup107, and if the N-terminal domains were independent and functioning correctly, the spNup131N+spNup132C protein should then be localized to the cytoplasmic side via Nup131N and the spNup132N+spNup131C protein should be retained on the nuclear side via Nup132N. The results showed that the spNup131N+spNup132C chimeric protein was predominantly diffused in the cytoplasm, with slight enrichment at the NE (**Fig 6B**) and that the other chimeric protein, spNup132N+spNup131C, also diffused to the cytoplasm but with no enrichment in the NE (**Fig 6B**). The relatively strong cytoplasmic staining of the chimeric proteins suggests that the N-terminal domains may function to prevent “abnormal” localization of spNup131 and spNup132. Overall, this result indicates that both N-terminal and C-terminal domains are required for the proper differential localization of spNup131 and spNup132. Consistent with this result, expression of these chimeric proteins failed to overcome the TBZ sensitivity in *nup132*Δ (**S7B Fig**), and to rescue the defects in meiosis in *nup132*Δ (**S7C Fig**).

**Fig 6.**
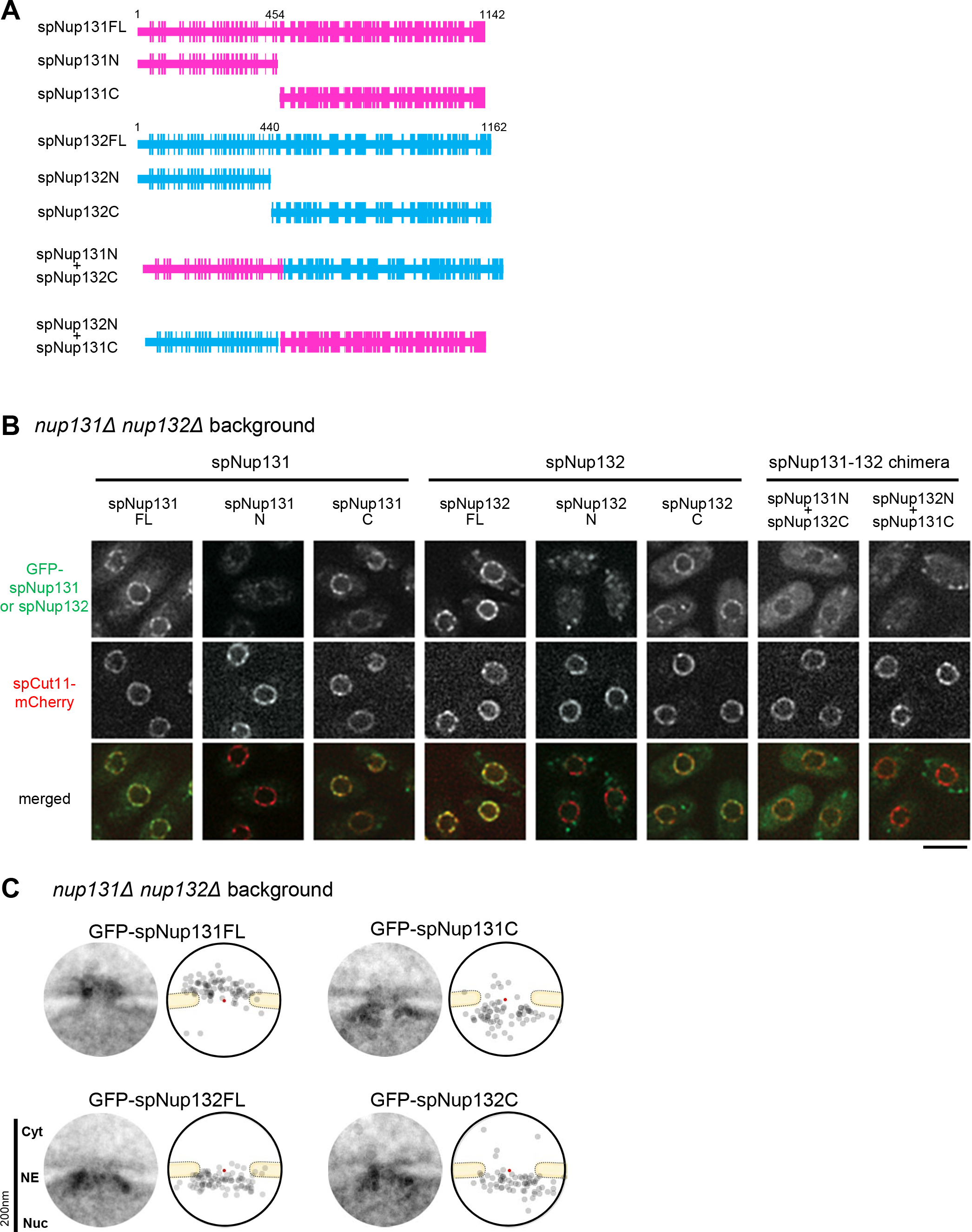
Localization of the spNup131 and spNup132 domains. **(A)** Schematics of the spNup131 and spNup132 fragments. Magenta represents peptides from spNup131 and light blue represents peptides from spNup132. **(B)** Localization of spNup131 and spNup132 fragments as determined by FM. GFP-fused fragments were expressed under the spNup132 promoter in the background of the *nup131*Δ *nup132*Δ double mutant. Cells were grown on YES plate media for only one day. For this short period of incubation, the NPC-clustering phenotype was not apparent. spCut11-mCherry served as an NPC marker. Single section images of the same focal plane are shown. Scale bar, 5 μm. **(C)** IEM of the spNup131 and spNup132 fragments. Projection images (left) and immunogold distributions (right) are shown as described in the legend of **Fig 1B**. The red points indicate the pore centers. IEM micrographs used for the quantitative analysis are available in S10 Dataset.

### Immunoelectron microscopy of the other nucleoporins

Because the Nup107-160 subcomplex seems to have a unique organization in *S. pombe*, we wished to understand the overall structure of the NPC. For this purpose, we performed IEM of other Nups to reveal their positionings within the NPC. GFP-fused Nups were used and IEM was carried out using a specific antibody against GFP, unless specified otherwise (**Fig 7**). We first examined inner ring Nups known as the Nup93 subcomplex in vertebrates. spNup97 and spNpp106, redundant *S. pombe* homologs of scNic96/hsNup93, were both similarly positioned near the center of the NPC (**Fig 7A**); the redundancy of scNic96/hsNup93 homologs is unique to the *Schizosaccharomyces* genus. spNup184 (scNup188/hsNup188) and spNup186 (scNup192/hsNup205) were also positioned near the center (**Fig 7A**). spNup40 (scNup53/scNup59/hsNup35) and spNup155 (scNup157/scNup170/hsNup155) were also located near the center of the NPC, but they showed a slightly broader range of localization (**Fig 7A**).

**Fig 7.**
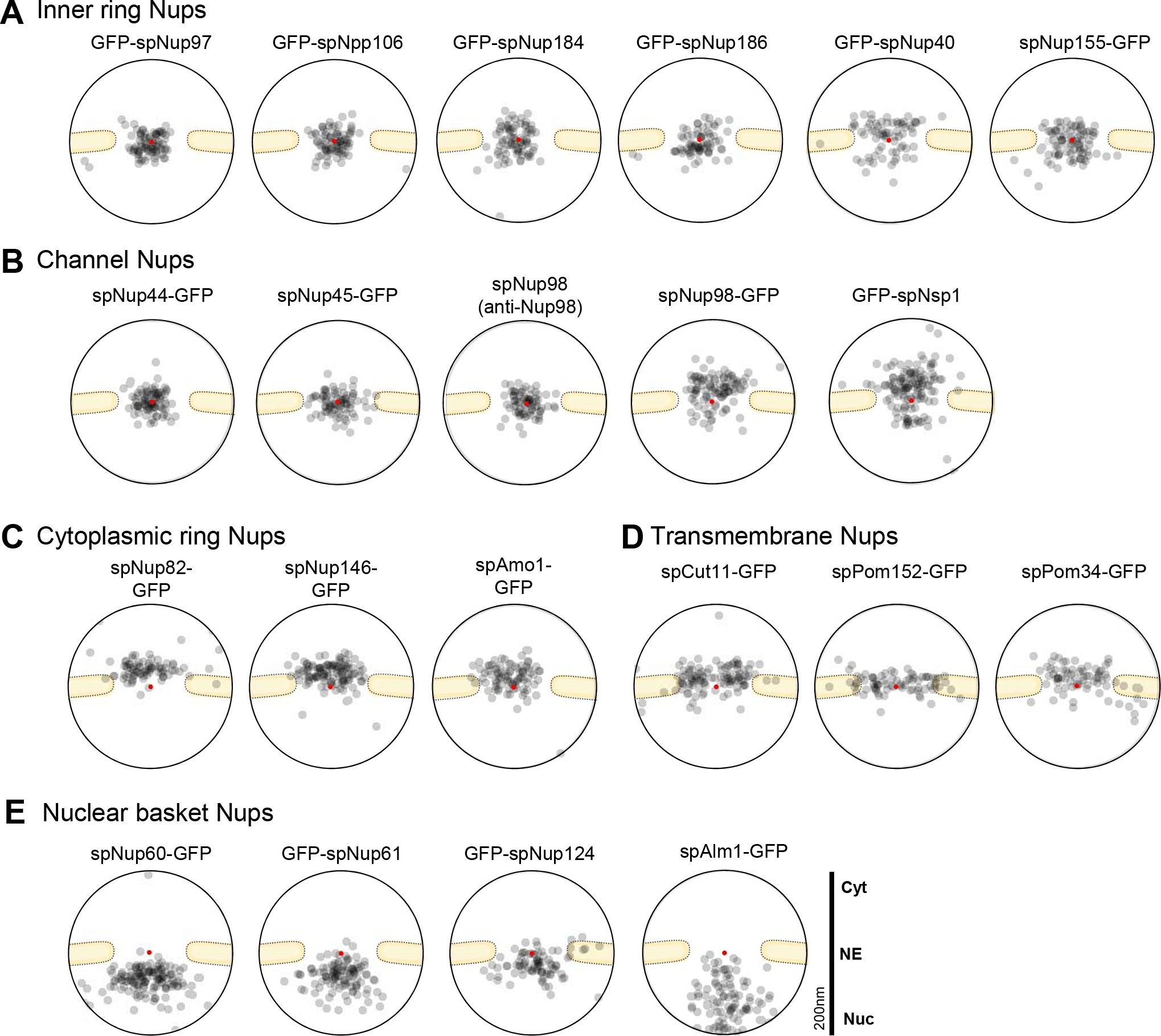
IEM of *S. pombe* nucleoporins. **(A)** Inner ring Nups. **(B)** Channel Nups. IEM for spNup98 was performed using an anti-Nup98 antibody (13C2) for the wild type strain and an anti-GFP antibody for the spNup98-GFP strain. **(C)** Cytoplasmic ring Nups. **(D)** Transmembrane Nups. **(E)** Nuclear basket Nups. The localization of the other nuclear basket Nup, spNup211, is shown in **Fig 3A**. Immunogold distributions are shown based on the projected immunoelectron micrographs for quantitative representation. The red points indicate the pore centers. Projections of raw IEM images are shown in S5C-G Fig. Individual IEM micrographs of 20 NPCs are available in S11 Dataset.

The channel Nups spNup44 (scNup57/hsNup54) and spNup45 (scNup49/hsNup58) were positioned at the center of the pore (**Fig 7B**). spNup98 (scNup145n/hsNup98) was detected at the center of the pore, when using an anti-Nup98 antibody (clone 13C2) [45], which specifically recognizes GLFG repeats located at the N-terminal region of endogenous spNup98 (see “spNup98” in **Fig 7B**). On the other hand, spNup98 (scNup145n/hsNup98) that was fused with GFP at its C-terminus was detected on the cytoplasmic side of the nuclear pore, when using an anti-GFP antibody (see “spNup98-GFP” in **Fig 7B**). In *H. sapiens* and *S. cerevisiae*, the C-terminal region of the Nup98 homologs (hsNup98/scNup145n/scNup100/scNup116) interacts with Nup96 homologs (hsNup96/scNup145c) [46–49], and the C-terminal Nup98-APD (autoproteolytic and NPC-targeting domain) binds to Nup88 homologs (hsNup88/scNup82) and to Nup96 in a mutually exclusive manner [49]. Thus, the C-terminal region of spNup98 is located near the positions of spNup96 and spNup82, both of which are positioned at the cytoplasmic side of the NPC, while the N-terminal region is extended to the center of the pore. The *S. pombe* homolog of the conserved Nup Nsp1 (scNsp1/hsNup62) was localized frequently in the cytoplasmic side and infrequently in the nuclear side of the NPC (**Fig 7B**).

spNup82 (scNup82/hsNup88) and spNup146 (scNup159/hsNup214) were detected at the cytoplasmic side (**Fig 7C**) as expected from their homologous counterparts. spAmo1(scNup42/hsNlp1) was also detected at the cytoplasmic side (**Fig 7C**). The transmembrane Nups, spCut11 (scNdc1/hsNdc1), spPom152 (scPom152), and spPom34 (scPom34) were localized at the center of the pore and slightly biased toward the cytoplasm (**Fig 7D**). The conserved Nups, spNup60 (scNup60), spNup61 (scNup2/hsNup50), and spNup124 (scNup1/hsNup153) (**Fig 7E)** as well as spNup211(scMlp1/scMlp2/hsTpr) (**Fig 3A**) were detected at the nuclear side as expected from their homologous counterparts. We added a recently identified Nup spAlm1 [50] to the group of nuclear Nups, according to its nucleoplasmic localization **(Fig 7E)**.

We summarized the positionings of the *S. pombe* Nups within the NPC in **Fig 8**. Transmembrane Nups (spCut11, spPom152, and spPom34); inner ring Nups (spNup97, spNpp106, spNup184, spNup186, spNup40, and spNup155); cytoplasmic ring Nups (spNup82, spNup146, spAmo1, spNsp1); central channel Nups (spNsp1, spNup98, spNup44, and spNup45); and nuclear basket Nups (spNup124, spNup60, spNup61, spNup211, and spAlm1) are positioned similarly to the positions of their orthologs in *S. cerevisiae* and human [1,4,5,26,51]. Thus, these NPC substructures, which include a central channel structure that is required for nucleocytoplasmic transport, seem to be common to other eukaryotes. In contrast, the outer ring Nup107-160 subcomplex (spNup131, spNup120, spNup85, spNup96, spNup37, spEly5 and spSeh1, spNup107 and spNup132) in *S. pombe* has a highly unusual asymmetrical organization: It splits into two pieces (spNup120 and spNup132 subcomplexes). The spNup120 subcomplex (spNup131, spNup120, spNup85, spNup96, spNup37, spEly5 and spSeh1) is located in the cytoplasmic side of the NPC while the spNup132 subcomplex (spNup107 and spNup132) is located in the nuclear side of the NPC. This asymmetrical organization of the Nup107-160 subcomplex may be required for building and maintaining the structural organization of the central channel and other NPC elements common to other eukaryotes, and consequently, the Nup107-160 subcomplex would be required for normal cell cycle progression in *S. pombe*.

**Fig 8.**
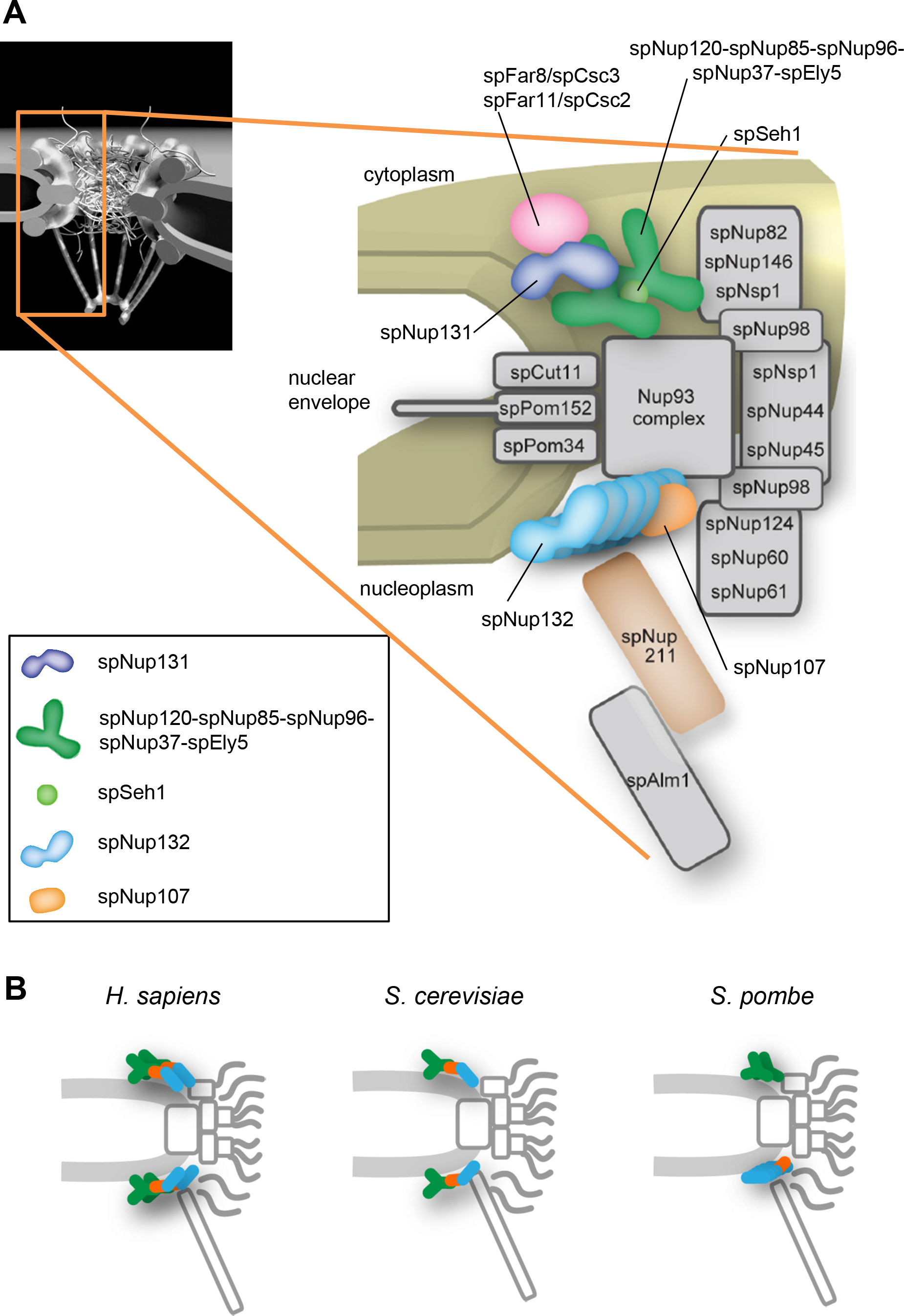
Model of the *S. pombe* NPC. **(A)** Positions of the *S. pombe* Nups. The numbers of the Nup107-160 subcomplex components were deduced based on the FM and biochemical analysis reported previously [29]: spNup131 (1 molecule, dark blue), the spNup120-spNup85-spNup96-spNup37-spEly5 complex (2 molecules each, dark green), spSeh1 (1 molecule, light green), spNup107 (1 molecule, orange), spNup132 (6 molecules, right blue). **(B)** Schematic drawings of the NPC structure in *H. sapiens* (left) [5] and *S. cerevisiae* (middle) [26] in comparison to that in *S. pombe* (right).

## Discussion

IEM of *S. pombe* Nups suggest that the *S. pombe*-specific Nup107-160 subcomplex structure is uniquely split into two pieces that localize differentially to the cytoplasmic and nuclear sides of the NPC while preserving the conserved modular structures of the Nup107-160 subcomplexes of other eukaryotes. High-precision distance measurements using FM also support this result (**Fig 1D, E, and 4b**). The asymmetrical organization of the *S. pombe* Nup107-160 subcomplex contrasts with the localization of the Nup107-160 subcomplexes in *H. sapiens*, *S. cerevisiae* and *Trypanosoma brucei*, in which Nup107-160 complexes are found on both the cytoplasmic and nuclear sides of the NPC [1,5,13,26,52]. Our results suggest that the *S. pombe* NPC has a novel organization that has evolved in the *Schizosaccharomyces* genus, which commonly bears the Nup132 and Nup131 clades (**S1B Fig**).

In *H. sapiens* and *S. cerevisiae*, the Nups in the Nup107-160 subcomplex assemble to form Y-shaped structures [5,21,24,25,53]. A total of 32 Y-complexes form two concentric reticulated rings at both the nuclear and cytoplasmic sides of the human NPC [4,5]. This organization may be supported by the iso-stoichiometry of each Nup in the Nup107-160 subcomplex: the amounts of each Nup in the Nup107-160 subcomplex in human cells are nearly equal [54]. In contrast, in *S. pombe*, the amounts of each Nup in the Nup107-160 subcomplex are not equal but highly divergent, ranging from 1 copy to 5-6 copies/unit [29]. In addition, the Sec13 homolog in *S. pombe* does not localize in the NPC, and Nup43 is not conserved in the genome [29]. The different stoichiometry and composition of the *S. pombe* Nup107-160 subcomplex components are likely due to the unique fission yeast–specific separated structure of the Nup107-160 subcomplex revealed in this study.

The mechanisms underlying the difference in the positioning of spNup131 and spNup132 at the NPC are still uncharacterized. Affinity capture/mass spectrometry showed that both spNup131 (cytoplasmic side) and spNup132 (nuclear side) bind to spNup107 (nuclear side) and spNup85 (cytoplasmic side). Considering that the molecular structural features of the spNup131 and spNup132 proteins are similar to each other, it is possible that both Nup131 and Nup132 interact with spNup107 and Nup85 *in vitro* in the whole cell extract, where there is no steric hindrance. In contrast, under *in vivo* conditions, the steric hindrance of these proteins can be a more important factor in determining their position in the NPC structure. In fact, neither spNup131N+spNup132C nor spNup132N+spNup131C was properly positioned in the NPC (Fig. 6B), supporting this notion. Although affinity capture/mass spectrometry showed the above-mentioned confusing results, this analysis also showed the interactor specific to each of spNup131 and spNup132. spNup146 (cytoplasmic side) appeared as a specific interactor to spNup131 (S3A Fig). On the other hand, spNup211 (nuclear side) appeared as a specific interactor to spNup132 (S3B Fig). These results suggest that spNup146 and spNup211 may be involved in the differential localization of spNup131 and spNup132, respectively. In fact, enforced relocalization of spNup107 to the cytoplasm side by expressing the spNup96-spNup107 fusion protein translocated half the fraction of spNup132 to the cytoplasmic side, while the other half remained in the nuclear side (Fig. 5C), suggesting that at least one more factor, other than spNup107, is involved in the positioning of spNup132. spNup211 can be such a factor that positions spNup132 in the nuclear side of the NPC.

Two Nup133 homologs spNup131 and spNup132 play different roles with different positions at the NPC. spNup132 is required for normal kinetochore formation during meiosis in *S. pombe* [30]. Deletion of the *nup132* gene but not the *nup131* gene causes untimely kinetochore assembly and activates spindle assembly checkpoint machinery during the first meiotic chromosome segregation [30]. The deletion of the *nup132* gene also increases the sensitivity to the microtubule-destabilizing drug thiabendazole, likely due to defects in the kinetochore structure in mitosis [29]. Similarly, in mammalian cells, kinetochore-related functions have been reported for the Nup107-160 subcomplex. A fraction of the Nup107-160 subcomplex is found at the kinetochores and spindle poles during mitosis [16,17,55,56], and depletion of the Nup107-160 subcomplex from kinetochores results in altered kinetochore protein recruitment [57–59]. While the molecular mechanism of underlying spNup132-mediated regulation of kinetochore proteins remains unknown, considering the similarity in kinetochore-related functions, spNup132, but not spNup131, is likely to be a functional homolog of mammalian Nup133.

On the other hand, spNup131 plays a different role at the cytoplasmic side of the NPC. IEM analysis in this study revealed that spNup131 is localized only on the cytoplasmic side of the NPC, and a genetic analysis showed interaction between spNup131 and spFar8. spFar8, which was proven to be located at the cytoplasmic side of the NPC in this study, is an *S. pombe* ortholog of the STRIPAK complex component Striatin [38]. In *S. pombe*, the STRIPAK complex regulates the septation initiation network through the conserved protein kinase Mob1 [37,60] and is required for asymmetric division of mother and daughter SPBs during mitosis [39]. The spNup131-dependent NPC localization of spFar8 revealed by this study implies that the NPC regulates STRIPAK localization in interphase cells. Human STRIPAK complexes have been proposed to play roles at the interface between the Golgi and the outer nuclear envelope [38]. Considering the physical and genetic interaction with spNup131, STRIPAK is likely to interact with the NPC on the cytoplasmic side of the NE. Although the role of the STRIPAK complex in interphase cells in *S. pombe* is not fully understood, the interaction between spNup131 and spFar8 may provide an important example linking the NPC to cytoplasmic structures.

This study demonstrates that the *S. pombe* Nup107-160 subcomplex has a novel separated structure and exhibits a localization pattern not reported in other organisms. Recent studies suggest that NPC structures are not necessarily the same among eukaryotes [26,61,62]. For example, the binucleated ciliate *Tetrahymena thermophila*, a single-cell organism, has two functionally distinct nuclei, the macronucleus (MAC) and micronucleus (MIC), in a single cell: The MAC and MIC differ in size, transcriptional activity, and nucleocytoplasmic transport specificity [63]. Interestingly, in *T. thermophila*, the MAC and the MIC NPCs are composed of differ amounts of the Nup107-160 subcomplex components [61]. The amount of the subcomplex in the MIC is about three times more than that in the MAC, suggesting that the Nup107-160 subcomplex forms different structures in the MAC from the MIC. In the green algae *Chlamydomonas reinhardtii*, a single cell organism, the NPCs have 24 Nup107-160 subcomplexes asymmetrically distributed within a single NPC: 16 at the nuclear side forming double outer rings and 8 at the cytoplasmic side forming one outer ring [62]. The asymmetrical distribution of the Nup107-160 subcomplex in *C. reinhardtii* may suggest a specific function of the complex in either cytoplasmic or nuclear side of the NPC although it is currently unclear. In some multicellular organisms, the expression level of Nups varies between cell types and during development [54,64–67]. It is also known that some mutations in Nups result in developmental defects in metazoans [68]. These findings suggest that the NPC composition in cell types and during development is biologically significant. Altered compositions might alter the NPC structure, at least in part; thus, unidentified NPC subcomplex structures may play roles in specific biological events such as differentiation and development. Thus, we speculate that novel NPC structures may be important to elucidate the biological functions of the NPC.

## Materials and Methods

### S. pombe cell cultivation

Strains used in this study are listed in S2 Table. YES or EMM2 culture medium was used for routine cultures [69]. For fluorescence microscopy, cells were grown in EMM2 liquid medium. ME medium was used to induce meiosis and spore formation. When necessary, TBZ (T5535-50G, Sigma-Aldrich Inc, Tokyo, Japan) was added to the YES medium to a final concentration of 20 μg/mL.

### Immunoelectron microscopy

For immunoelectron microscopy, 1.5 × 10^8^ cells expressing GFP-Nup131 were fixed in 1 mL of a mixture of 4% formaldehyde (18814-10, Polysciences, Inc, Warrington, PA, USA) and 0.01% glutaraldehyde (1909-10, Polysciences, Inc.) dissolved in 0.1 M phosphate buffer (PB) (pH 7.4) for 20 min at room temperature, treated with 0.5 mg/mL Zymolyase 100T (7665-55, Nacalai Tesque, Inc., Kyoto, Japan) in PB for 20-30 min at 30°C, and then permeabilized with 0.2% saponin (30502-42, Nacalai Tesque, Inc.) and 1% bovine serum albumin (BSA) in PB for 15 min. The GFP epitope tag was labeled with a primary antibody (600-401-215, rabbit polyclonal anti-GFP antibody, Rockland Immunochemicals, Limerick, PA, USA) diluted 1:400 in PB containing 1% BSA and 0.01% saponin, and a secondary antibody (7304, goat anti-rabbit Alexa 594 FluoroNanogold Fab’ fragment; Nanoprobes Inc., Yaphank, NY, USA) diluted 1:400. The same immunostaining conditions were used for the cells expressing each of the other GFP-fused Nups except for the dilution ratios of primary and secondary antibodies; the conditions used for each experiment are listed in S3 Table. For analysis of the spNup98 N-terminal region, we used a mouse monoclonal anti-Nup98 antibody (13C2) [43,45] (available from BioAcademia Inc, Japan, Cat. #70-345) diluted 1:100 and anti-mouse Alexa594 FluoroNanogold Fab’ fragment (7302, Nanoprobes) diluted 1:400. Cells then were fixed again with 1% glutaraldehyde in PB for 1 h at room temperature and treated with 100 mM lysine HCl in PB twice for 10 min each. The cells were stored at 4°C until use. Before use, the cells were incubated with 50 mM HEPES (pH 5.8) three times for 3 min each and with distilled water (DW) once, incubated with Silver enhancement reagent (a mixture of equal volumes of the following A, B, and C solutions: A, 0.2% silver acetate solution; B, 2.8% trisodium citrate-2H_2_O, 3% citric acid-H_2_O, and 0.5% hydroquinone; C, 300 mM HEPES, pH 8.2) at 25°C for 2-5 min. Cells were embedded in 2% low melting agarose dissolved in DW. Cells were post-fixed with 2% OsO_4_ in DW for 15 min and stained with 1% uranyl acetate in DW at room temperature. Cells were dehydrated using stepwise incubations in ethanol and acetone and finally embedded in epoxy resin Epon812. Solidified blocks containing cells were sectioned, and the ultra-thin sections were stained with uranyl acetate and lead citrate, the usual pretreatment for EM observations. Images were obtained using a JEM1400 transmission electron microscope (JEOL, Tokyo, Japan) at 120kV.

Nuclear pores containing more than two immunogold particles were chosen for localization analysis as described previously [1]. To confirm the accessibility of the nucleus to immunogold particles, the nuclear centromere protein spMis6-GFP was co-expressed with GFP-fused Nups in cells and stained with anti-GFP antibody for IEM. For quantification of the Nup signals, we chose only cell specimens with a positive spMis6-GFP signal. For quantitative representations, montage pictures were produced by stacking 20 NPC images with 5% opacity on Adobe Photoshop CS software.

We also quantified the distribution of GFP-fused proteins in the cytoplasmic, middle, and nucleoplasmic regions of the NPC by counting the number of gold particles in each region (**S8 Fig**).

### cDNA clones

*nup107*^+^ and *nup37*^+^ cDNAs were provided by National BioResource Project Japan (http://yeast.lab.nig.ac.jp/yeast/top.xhtml). cDNA fragments of other Nups were amplified from a cDNA library pTN-RC5 or pTN-FC9 (National BioResource Project Japan) using PCR.

### Strain construction

To visualize nuclear pore localization of the different domains of Nup131 and Nup132, *lys1*^+^-integrating plasmids carrying GFPs65t-fused with respective Nup domains were introduced into cells with a *nup131*Δ*nup132*Δ double-deletion mutant background. To construct the spNup96-spNup107 fusion Nup, a cDNA fragment encoding spNup107 or spNup107-GFP followed by a drug resistance marker gene was integrated after the chromosomal spNup96 coding region. After a diploid strain was obtained by crossing the spNup96-spNup107 fusion containing strain with the wild type strain, the *nup107*^+^ gene on the original chromosomal locus was deleted. To introduce the chromosomal fluorescent tag and gene disruption, a two-step PCR method was applied [70]. Nup-GFP fusion constructs were described previously [29]. The spMis6-GFP fusion was constructed as described previously [71] or using the two-step PCR method. mCherry-spAtb2 was visualized as described previously [30].

### Plasmid construction

To express the full-length or domains of the spNup131 and spNup132 proteins, cDNA fragments were amplified by PCR. For overexpression, the PCR products were sub-cloned into the plasmid that carries the *nmt1* promoter and terminator. For physiological expression, the PCR products were sub-cloned into the *Bgl*II site of the plasmid (pY53) that carries the *nup132* promoter (−1000bp)-driven GFPs65t and the *nmt1* terminator. Each sub-cloning was done by using the In-Fusion PCR cloning kit (Clontech Laboratories, Mountain View, CA, USA). Plasmids were integrated at the *lys1* gene locus. Correct integrations were confirmed by PCR.

### Fluorescence microscopy

Images were obtained using a DeltaVision microscope system (GE Healthcare, Tokyo, Japan) equipped with a CoolSNAP HQ^2^ CCD camera (Photometrics, Tucson, AZ, USA) through an oil-immersion objective lens (PlanApoN60×OSC; NA, 1.4) (Olympus, Tokyo, Japan) as described previously [29]. Z-stack images were obtained and processed by deconvolution and denoising algorithm [72] when necessary. For time lapse microscopy, cells were observed every 5 minutes as described previously [30]. The projection images of z-stacks were made by softWoRx software equipped in the microscope system.

### Precise distance measurements with fluorescence microscopy

Cells to be analyzed were cultured in 150 μl of the EMM2 5S medium on an 8-well chambered coverglass, Lab-Tek II (Thermo Fisher Scientific Japan, Yokohama, Japan); cells used for references for chromatic correction were also cultured in a chamber of the same coverglass. Simultaneously acquired multicolor images were obtained with OMX SR (GE Healthcare) equipped with three sCMOS cameras. The three-dimensional (3D) images were deconvolved by constrained iterative deconvolution using the Priism suite (http://msg.ucsf.edu/IVE/) with a Wiener filter enhancement of 0.9 and 15 iterations. To correct for chromatic shifts of multicolor FM images, green-to-red photoconversion of GFP was used as the reference as previously reported [73,74]: Nup96-GFP was used in this study. For photoconversion, cells were illuminated with 405 nm light at about 393 W/cm^2^ for 4 seconds, then both green and red-converted GFP species were excited with 488 nm to obtain images of the same object in the green and red channels. Chromatic shifts were measured using such images, and the correction parameters were determined using the Chromagnon software (https://github.com/macronucleus/Chromagnon), by which a chromatic shift can be corrected with an accuracy of ~10 nm in 2D XY plane and ~15 nm in 3D XYZ space using test samples [35]. The correction parameters were applied to all images obtained from the same chambered coverglass as described in [35].

Next, individual nuclei in the images were identified and segmented by our software using functions from the “ndimage” module of the Scipy package (http://www.scipy.org). The centers of the nuclei were identified by least-square fitting with 3D ellipses using the 3D coordinates of the nucleoporin fluorescence above threshold. The images of the NE at the midsection were linearized with polar transformation. These images were then averaged across all angles to produce 1D intensity profiles along the radial direction of the nuclei. The intensity profiles were fitted to Gaussian profiles to determine a distance between the peaks of green and red for each of the individual nuclei. The mean distance was determined from more than 70 nuclei. Because not all nuclei were round, we rejected nuclei of elliptical or irregular shapes before the final calculation.

### Western blot analysis

For western blot analysis, whole cell extracts were prepared from approximately 5×10^6^ cells by trichloroacetic acid (TCA) precipitation as described previously [29]. To detect GFP-fused proteins, a rabbit polyclonal anti-GFP antibody (600-401-215, Rockland Immunochemicals Inc.) was used. To detect spNup98, a mouse monoclonal anti-Nup98 antibody (13C2) was used [45]. HRP-conjugated goat anti-rabbit or anti-mouse IgG (NA9340-1ml or NA9310-1ml, GE Healthcare) was used as a secondary antibody. Protein bands were detected by chemiluminescence using ChemiDoc MP imaging system (Bio-Rad).

### Preparation of whole cell extracts for affinity-capture experiments

Growing cells (about 5 × 10^9^) were collected and washed with 10 mM HEPES buffer (pH 7.5) containing 1 mM phenylmethylsulfonyl fluoride (PMSF). The washed cells were divided into aliquots of 3 × 10^8^ cells in 10 mM HEPES buffer (pH 7.5) containing 1 mM PMSF. Cells were again collected by centrifugation, and the cell pellet was kept frozen by liquid nitrogen until use. To make a cell extract, the cell pellet was thawed and suspended in 100 μL of lysis buffer (50 mM HEPES (pH 7.5), 150 mM NaCl, 1% Triton X-100, 1 mM EDTA, 2 mM PMSF) with a protease inhibitor cocktail (165-20281, Wako, Tokyo, Japan) and mashed by glass beads using Multi-beads shocker (Yasui Kikai Corporation, Osaka, Japan). We chose this extraction condition based on a previous study identifying the components of plant NPCs [11]. Because the presence of 1% Triton X-100 gave better results for extracting plant NPC components, we decided to use 1% Triton X-100 as a detergent, and we examined three different salt conditions (50 mM, 150 mM or 500 mM NaCl) in the presence of the detergent. As a result, 150 mM NaCl gave better results with low background. At least for extracting spNup131 and spNup132, this condition was better than that optimized for extracting NPCs in other organisms such as budding yeast (1.5 M ammonium acetate, 1% Triton X-100) [75] and trypanosoma (20 mM HEPES (pH7.4), 250 mM NaCitrate, 0.5% Triton X-100, 0.5% deoxy Big CHAP) [76]. After further addition of 400 μL of lysis buffer, the mashed cell pellet was transferred to new microtubes. The supernatant was collected after centrifugation at 15000 rpm for 15 min at 4°C and used as the whole-cell extract.

### Affinity-capture and LC/MS/MS analysis

The whole-cell extract was incubated with a rabbit anti-GFP antibody (600-401-215, Rockland Immunochemicals). Antibody-conjugated proteins were collected by incubating with Protein A Sepharose beads (17528001, GE Healthcare). Beads were then washed 4-5 times with the lysis buffer described above. After elution in SDS-PAGE sample buffer, protein samples were loaded onto a 12% SDS-PAGE gel for liquid chromatography coupled to tandem MS (LC/MS/MS). Data analysis for LC/MS/MS was performed as described previously [77] using the Pombase protein dataset released on November 12^th^, 2015. To identify proteins interacting with spNup131 and spNup132, protein samples were prepared from two independent experiments and each preparation was analyzed by LC/MS/MS. Proteins detected as more than one unique spectrum were identified as interacting proteins with spNup131 and spNup132. Alternatively, when proteins detected as only one unique spectrum in the first experiment were repeatedly detected in the second experiment, they were also identified as interacting proteins.

## Acknowledgments

We thank Naomi Takagi for technical assistance for mass spectrometry analysis and the National BioResource Project Japan for the Nup cDNA clones. We also thank Drs. Thomas U. Schwartz, Valérie Doye, and David B Alexander for critical reading of this paper.

## Supporting information

**S1 Fig.** Phylogenetic analysis of Nup133 proteins

**S2 Fig.** IEM images of spMis6-GFP, GFP-spNup131, and GFP-spNup132

**S3 Fig.** Affinity capture/mass spectrometry of GFP-spNup131 and GFP-spNup132

**S4 Fig.** FM images of spFar11-GFP in wild type, *nup131*Δ, and *nup132*Δ cells

**S5 Fig.** Projections of raw IEM images

**S6 Fig.** Duration of meiosis I and II in *nup131*Δ cells

**S7 Fig.** Characterization of the strains used in Fig. 6

**S8 Fig.** Distribution of Nups within the NPC

**S1 Table.** Nucleoporins in *S. pombe*, *S. cerevisiae*, and *H. sapiens*

**S2 Table.** *S. pombe* strains used in this study

**S3 Table.** Dilution ratios of primary and secondary antibodies used for IEM

**S1 Dataset.** Individual IEM images of 20 NPCs used for superimposed images of Fig 1C (spNup131-GFP and spNup132-GFP)

**S2 Dataset.** Values of the distance between mCherry-spNup132 and GFP-spNup131 and those between mCherry-spNup131 and GFP-spNup132 measured for Fig 1E

**S3 Dataset.** Individual IEM images of 20 NPCs used for superimposed images of Fig 2B (spFar8-GFP)

**S4 Dataset.** Individual IEM images of 20 NPCs and the projection image analyzed for Fig 3A (spNup211-GFP)

**S5 Dataset.** Values of the maximum fluorescence intensity of spNup211-GFP in wild type, *nup131*Δ and *nup132*Δ cells measured for Fig 3C

**S6 Dataset.** Individual IEM images of 20 NPCs and their projection images analyzed for Fig 4A (pNup120-GFP, spNup85-GFP, spNup96-GFP, spNup37-GFP, spEly5-GFP, spSeh1-GFP, spNup107-GFP and GFP-spNup107)

**S7 Dataset.** Values of the distance between spNup85-GFP and mCherry-spNup131 and those between spNup85-GFP and mCherry-spNup132 measured for Fig 4B

**S8 Dataset.** Values of the distance between spNup107-GFP and mCherry-spNup131 and those between spNup107-GFP and mCherry-spNup132 measured for Fig 4C

**S9 Dataset.** Individual IEM images of 20 NPCs and their projection images analyzed for Fig 5B (spNup96-spNup107-GFP) and 5C (GFP-spNup132 in the presence of the spNup96-spNup107 fusion protein)

**S10 Dataset.** Individual IEM images of 20 NPCs and their projection images analyzed for Fig 6C (full length GFP-spNup131 and GFP-spNup132, and their C-terminal domains in the *nup131*Δ*nup132*Δ mutant cells)

**S11 Dataset.** Individual IEM images of 20 NPCs and their projection images analyzed for Fig 7 (inner ring Nups, channel Nups, cytoplasmic ring Nups, transmembrane Nups and nuclear basket Nups)

