## Supplemental Figures and Tables for "Asymmetrical localization of Nup107-160 subcomplex components within the nuclear pore complex in fission yeast"

**Supporting information**

**S1 Fig.** Phylogenetic analysis of Nup133 proteins

**S2 Fig.** IEM images of spMis6-GFP, GFP-spNup131, and GFP-spNup132

**S3 Fig.** Affinity capture/mass spectrometry of GFP-spNup131 and GFP-spNup132

**S4 Fig.** FM images of spFar11-GFP in wild type, *nup131*Δ, and *nup132*Δ cells

**S5 Fig.** Projections of raw IEM images

**S6 Fig.** Duration of meiosis I and II in *nup131*Δ cells

**S7 Fig.** Characterization of the strains used in Fig. 6

**S8 Fig.** Distribution of Nups within the NPC

**S1 Table.** Nucleoporins in *S. pombe*, *S. cerevisiae*, and *H. sapiens*

**S2 Table.** *S. pombe* strains used in this study

**S3 Table.** Dilution ratios of primary and secondary antibodies used for IEM

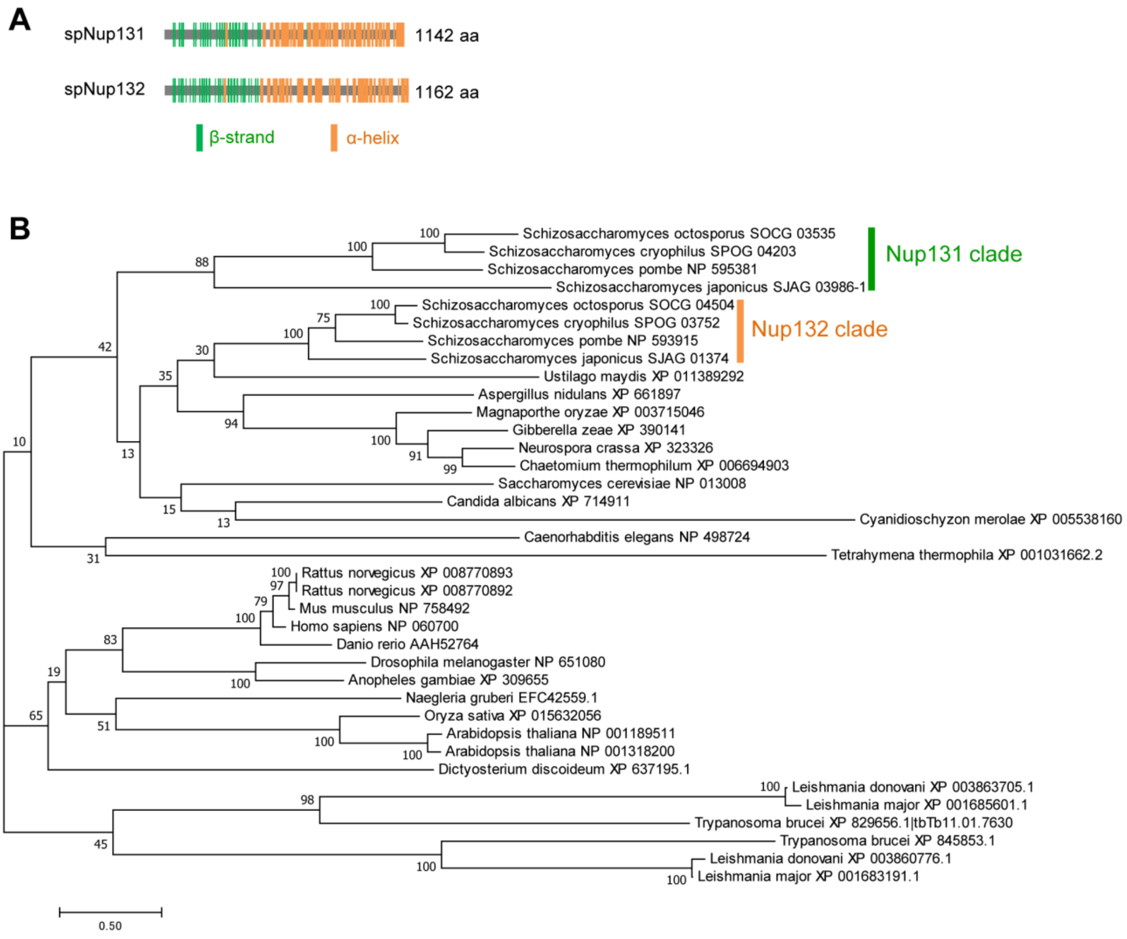


**S1 Fig. Phylogenetic tree of Nup133-like proteins**

**(A)** Distribution of secondary structure elements on spNup131 and spNup132. Both spNup131 and spNup132 have structural features in common with Nup133 found in many organisms, such as the N-terminal β-propeller rich region assigned as the Nup133 N-terminal like domain (Pfam PF08801; amino acid residues 44–454 in spNup131 and 50–440 in spNup132) and the C-terminal α-helical stack region assigned as the Non-repetitive/WGA-negative nucleoporin C-terminal domain (Pfam PF03177; a.a. residues 581–1049 in spNup131 and 515–1084 in spNup132). **(B)** Species names and Genbank accession numbers are shown. spNup131- and spNup132-like proteins found in fission yeasts are indicated. The sequences were aligned using Muscle in MEGA7 software. The evolutionary history was inferred by using the Maximum Likelihood method (Le and Gascuel, *Mol. Biol. Evol.* 25, 1307-1320, 2008). The tree with the highest log likelihood (-26923.30) is shown. The bootstrap values are presented next to the branches. Evolutionary analyses were conducted in MEGA7 (Kumar S et al., *Mol. Biol. Evol*. 33, 1870-1874, 2016).

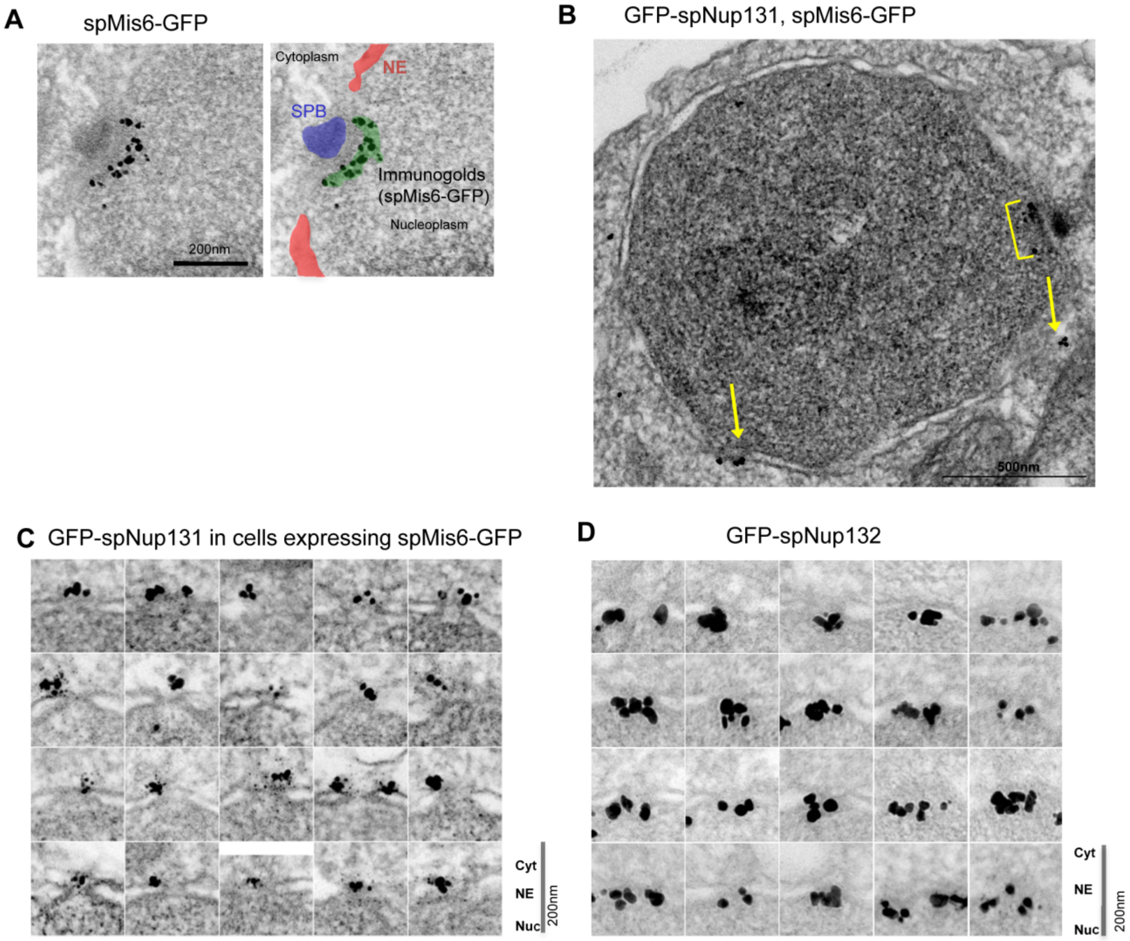


**S2 Fig. IEM images of spMis6-GFP, GFP-spNup131, and GFP-spNup132**
**(A)** IEM of spMis6-GFP. An original electron micrograph (left) and its duplicated image (right) indicating subcellular structures are shown. SPB, spindle pole body; NE, nuclear envelope. **(B)** IEM of co-expressed GFP-spNup131 and spMis6-GFP. A representative image is shown. Arrows indicate immunogold at the nuclear pores. The yellow-lined regions indicate immunogold near the SPB, corresponding to the signals from spMis6-GFP. **(C, D)** Immunoelectron micrographs of 20 nuclear pores used to generate the montage picture and distribution analysis in **Fig 1C**. Scale bars, 200 nm. **(C)** IEM of GFP-spNup131 and spMis6-GFP. **(D)** IEM of GFP-spNup132.

**A**

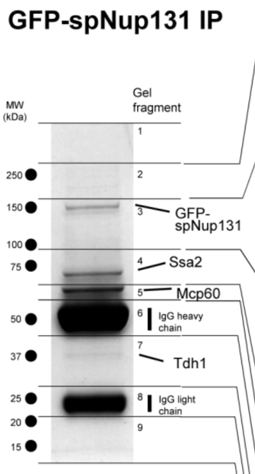


| Gel | Name | Accession No. | MW | Unique spectra |
|---|---|---|---|---|
| 2 | Utp20 | SPBC56F2.04 | 285.2 | 2 |
| | Tor2 | SPBC216.07c | 266.4 | 3 |
| | Cut6 | SPAC56E4.04c | 256.8 | 20 |
| | Mor2 | SPBP19A11.04c | 251.2 | 1 |
| | Ura1 | SPAC22G7.06c | 248.3 | 22 |
| | Not1 | SPAC20G8.06 | 237.1 | 6 |
| | Fas1 | SPAC926.09c | 230.6 | 43 |
| | Spo15 | SPAC1F3.06c | 222.8 | 3 |
| | Sec72 | SPAC30.01c | 206.8 | 2 |
| | Fas2 | SPAC4A8.11c | 202.2 | 47 |
| | Nuc1 | SPBC4C3.05c | 189.2 | 2 |
| | Myo2 | SPCC645.05c | 176.4 | 58 |
| | Myo52 | SPCC1919.10c | 175.2 | 3 |
| | Tif471 | SPAC17C9.03 | 154 | 2 |
| | Nup146 | SPAC23D3.06c | 145.8 | 6 |
| 3 | Nup131 | SPBP35G2.06c | 131.4* | 59 |
| | Trm72 | SPCC1494.07 | 171.2 | 2 |
| | Nup155 | SPAC890.06 | 147.6 | 2 |
| | Elp1 | SPBC36.07 | 141.6 | 2 |
| | Cop1 | SPBPJ4664.04 | 136.4 | 3 |
| | Rpa2 | SPBP23A10.07 | 131.7 | 1 |
| | Tea3 | SPAC6G10.02c | 127.8 | 2 |
| | Elf1 | SPAC3C7.08c | 116.7 | 3 |
| | Tps3 | SPACUNK4.16c | 106.8 | 2 |
| | Atg11 | SPAC7D4.04 | 106.8 | 2 |
| | Sec21 | SPAC57A7.10c | 100.9 | 4 |
| | Utp21 | SPCC1672.07 | 100.6 | 2 |
| | Def1 | SPBC354.10 | 100.1 | 1 |
| | ribosome biogenesis factor recycling AAA family ATPase | SPBC56F2.07c | 88.5 | 2 |
| 4 | Nup107 | SPBC428.01c | 93.3 | 19 |
| | Sec27 | SPBC16C6.13c | 89.8 | 1 |
| | Gap1 | SPBC646.12c | 87.5 | 1 |
| | Cdr2 | SPAC57A10.02 | 86 | 2 |
| | Ade1 | SPBC405.01 | 85.2 | 2 |
| | Sec231 | SPCC31H12.07 | 84.8 | 2 |
| | Leu2 | SPAC9E9.03 | 82.8 | 2 |
| | Nup85 | SPBC17G9.04c | 78 | 2 |
| | Ssc1 | SPAC664.11 | 73 | 28 |
| | Ssa2 | SPCC1739.13 | 70.2 | 58 |
| | Ssa1 | SPAC13G7.02c | 70.1 | 21 |
| | Far8 | SPBC1773.01 | 69.5 | 2 |
| | Dbp2 | SPBP8B7.16c | 61.5 | 2 |
| | Ght8 | SPCC548.06c | 60.2 | 1 |
| 5 | Acl1 | SPBC1703.07 | 67.2 | 1 |
| | Tnr3 | SPAC6F12.05c | 64.2 | 5 |
| | Mcp60 | SPAC12G12.04 | 62.2 | 82 |
| | Elp3 | SPAC29A4.20 | 61.8 | 2 |
| | Ade6 | SPCC1322.13 | 60 | 4 |
| | Gua1 | SPBC2F12.14c | 57 | 4 |
| 6 | Rvb1 | SPAPB8E5.09 | 50.1 | 3 |
| | Rpt6 | SPBC23G7.12c | 45.1 | 3 |
| 7 | Mpg1 | SPCC1906.01 | 39.7 | 1 |
| | Fba1 | SPBC19C2.07 | 39.6 | 1 |
| | Tdh1 | SPBC32F12.11 | 35.9 | 19 |
| 8 | Sno1 | SPAC222.08c | 25.9 | 1 |
| 9 | Dop1 | SPAPB21F2.02 | 192.7 | 1 |

**B**

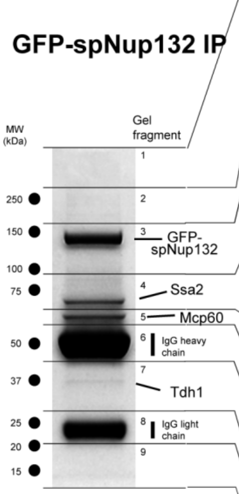


| Gel | Name | Accession No. | MW | Unique spectra |
|---|---|---|---|---|
| 1 | Mdn1 | SPCC737.08 | 537.8 | 13 |
| 2 | Brr2 | SPAC9.03c | 248.8 | 1 |
| | Ura1 | SPAC22G7.06c | 248.3 | 21 |
| | Fas1 | SPAC926.09c | 230.6 | 24 |
| | Nup211 | SPCC162.08c | 211.5 | 1 |
| | Fas2 | SPAC4A8.11c | 202.2 | 36 |
| | Myo2 | SPCC645.05c | 176.4 | 44 |
| 3 | Nup132 | SPAC1805.04 | 132.1* | 138 |
| | Nup155 | SPAC890.06 | 147.6 | 3 |
| | Rpa2 | SPBP23A10.07 | 131.7 | 1 |
| | Nup120 | SPBC3B9.16c | 129.8 | 3 |
| | Elf1 | SPAC3C7.08c | 116.7 | 1 |
| | Pfk1 | SPBC16H5.02 | 102.6 | 20 |
| | Eft201, Eft202 | SPAC513.01c, SPCP31B10.07 | 93.2 | 22 |
| 4 | Dop1 | SPAPB21F2.02 | 192.7 | 1 |
| | Kap95 | SPAC1B1.03c | 94.7 | 3 |
| | Nup96 | SPAC1486.05.2 | 94.6 | 1 |
| | Nup107 | SPBC428.01c | 93.3 | 17 |
| | Cdc22 | SPAC1F7.05 | 92 | 2 |
| | Sec231 | SPCC31H12.07 | 84.8 | 1 |
| | Nup85 | SPBC17G9.04c | 78 | 2 |
| | Ssc1 | SPAC664.11 | 73 | 23 |
| | Ssa2 | SPCC1739.13 | 70.2 | 66 |
| | Ssa1 | SPAC13G7.02c | 70.1 | 25 |
| | Sks2 | SPBC1709.05 | 67.2 | 20 |
| 5 | Mcp60 | SPAC12G12.04 | 62.2 | 68 |
| | Cut15 | SPCC962.03c | 60.4 | 2 |
| | Imp1 | SPBC1604.08c | 59.9 | 4 |
| | Ade4 | SPAC4D7.08c | 59.8 | 2 |
| | DNAJ domain protein | SPAC4H3.01 | 44.8 | 2 |
| 6 | Tcb3 | SPAPYUK71.03c | 135.8 | 1 |
| | Tef101 | SPCC794.09c | 49.7 | 23 |
| | Eno101 | SPBC1815.01 | 47.4 | 1 |
| 7 | Arp2 | SPAC11H11.06 | 44.2 | 1 |
| | Sam1 | SPBC14F5.05c | 41.8 | 1 |
| | Act1 | SPBC32H8.12c | 41.8 | 20 |
| | Sum1 | SPAC4D7.05 | 36.8 | 1 |
| | Tdh1 | SPBC32F12.11 | 35.9 | 24 |
| 8 | Rpl701 | SPBC18H10.12c | 28.7 | 1 |
| 9 | Cay1 | SPBC2F12.12c | 60.9 | 1 |
| | Sar1 | SPBC31F10.06c | 21.3 | 1 |

**C**

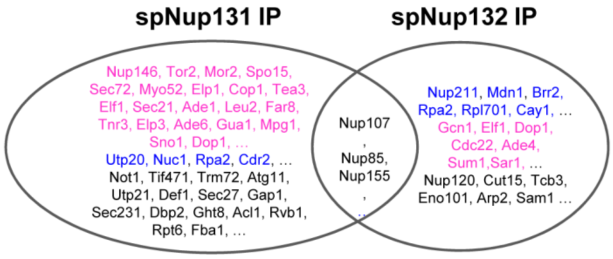


magenta, cytoplasmic proteins
Blue, nuclear proteins
Black, proteins of the other or unidentified localization

**S3 Fig. Affinity capture/mass spectrometry of GFP-spNup131 and GFP-spNup132**
**(A, B)** Proteins bound to GFP-spNup131 and GFP-spNup132. Images of Coomassie-stained SDS-PAGE gels are shown. Dots indicate the positions of molecular weight marker proteins shown on the left. Each gel was cut at the positions shown by the horizontal lines on the gel image. Proteins that correspond to major bands in each

gel fragment were deduced by LC/MS/MS analysis and are indicated on the right. The list on the right shows proteins specifically bound to GFP-spNup131 and GFP-spNup132, Nups, and abundant proteins (>20 spectra). Protein names are colored by their subcellular localizations according to gene ontology data (Pombase: https://www.pombase.org/): magenta, cytoplasmic proteins; blue, nuclear proteins; black, proteins of other or unidentified localizations. **(C)** Venn diagram showing proteins bound to GFP-spNup131 and GFP-spNup132 identified by LC/MS/MS analysis. Protein names are colored by their subcellular localizations: magenta, cytoplasmic proteins; blue, nuclear proteins; black, proteins of other or unidentified localizations.

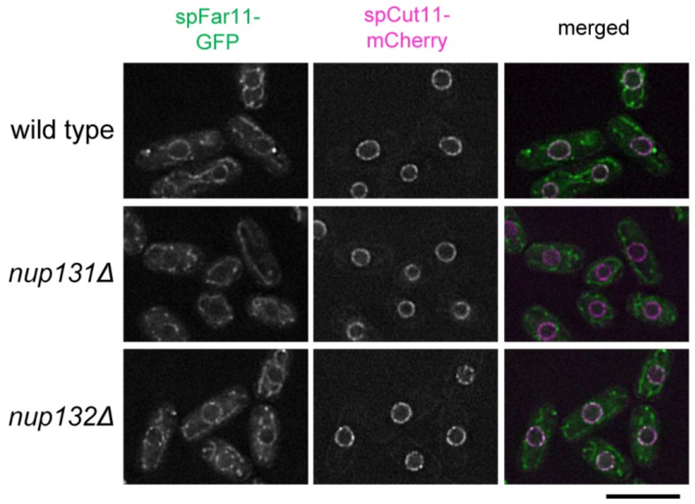


**S4 Fig. FM images of spFar11-GFP in wild type, *nup131*Δ, and *nup132*Δ cells**
Cells were prepared and observed as described in **Fig 2A**. Scale bar, 10 μm.

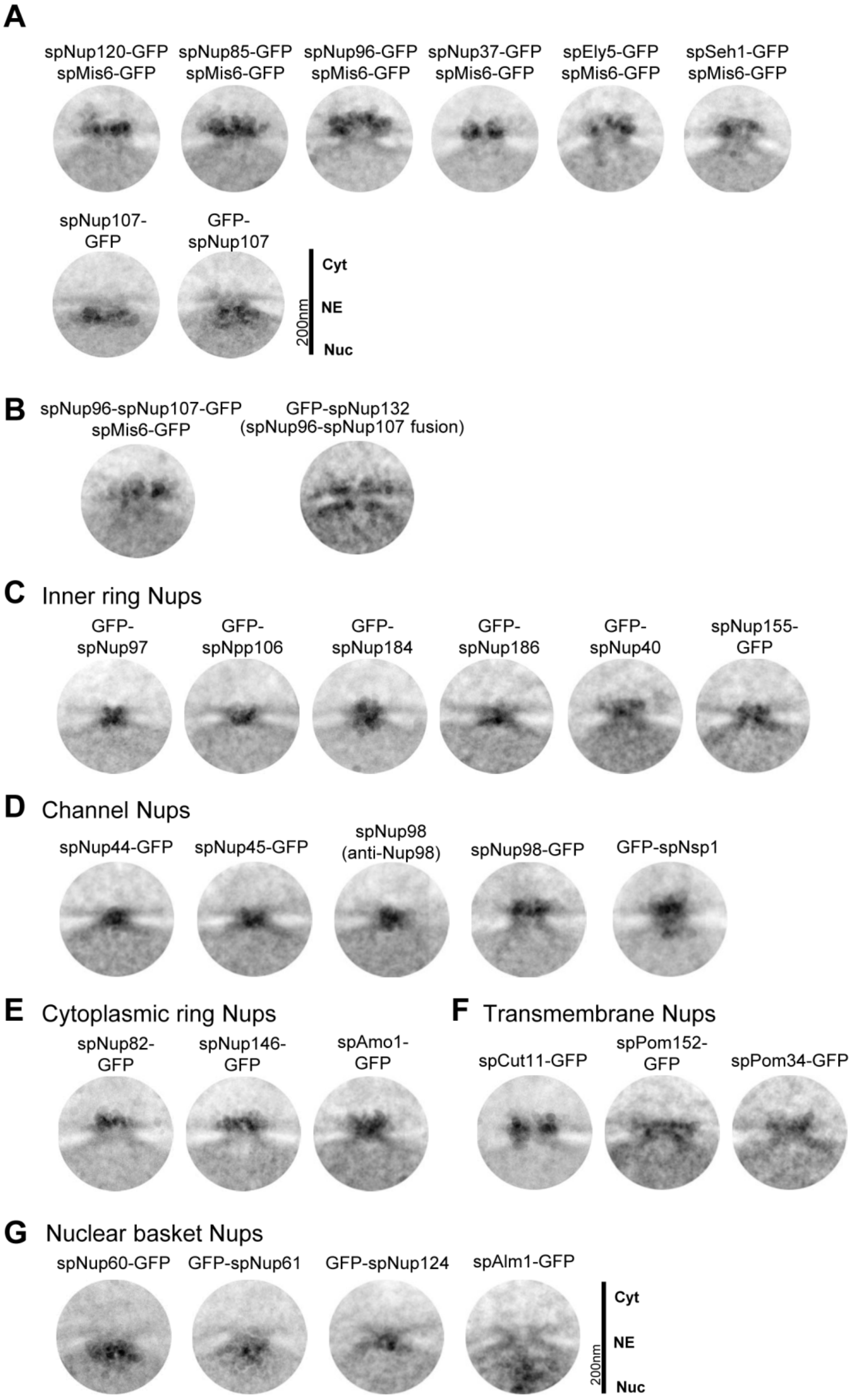


**S5 Fig. Projection of raw IEM images**

Projection images were made from 20 NPC IEM images for each Nups. **(A)** Outer ring Nups. Individual IEM images are available in S6 Dataset. **(B)** spNup96-spNup107-GFP fusion protein and GFP-spNup132 in spNup96-spNup107 fusion strain. Individual IEM images are available in S9 Dataset. **(C)** Inner ring Nups. **(D)** Channel Nups. **(E)** Cytoplasmic ring Nups. **(F)** Transmembrane Nups. **(G)** Nuclear basket Nups. Individual IEM images for (C)-(G) are available in S11 Dataset.

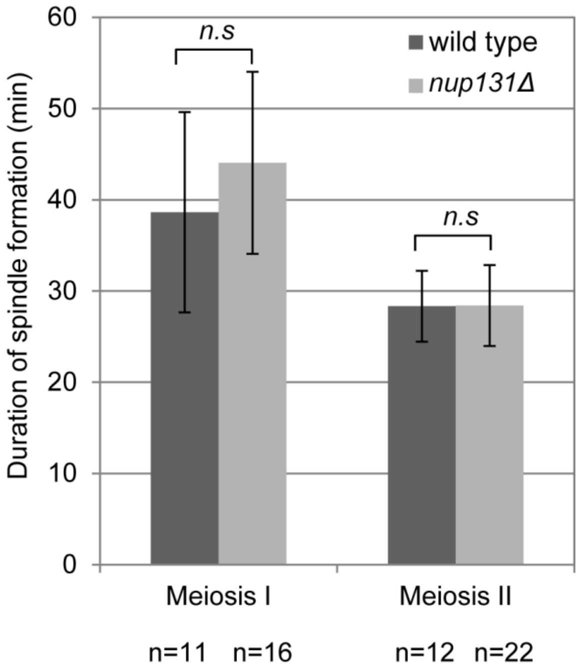


**S6 Fig. Durations of meiosis I and II in *nup131*Δ cells**

Durations of meiosis I and meiosis II were measured by time-lapse observation. Error bars represent standard deviations. The duration of meiosis I was 38.6 ± 11.0 min in wild type and 44.1 ± 10.0 min in *nup131*Δ cells ($p$=0.41, student's t-test); the duration of meiosis II was 28.3 ± 3.9 min in wild type and 28.4 ± 4.4 min in *nup131*Δ cells ($p$=0.96, student's t-test). n.s. stands for no significant difference. Numbers of observed cells are indicated at the bottom.

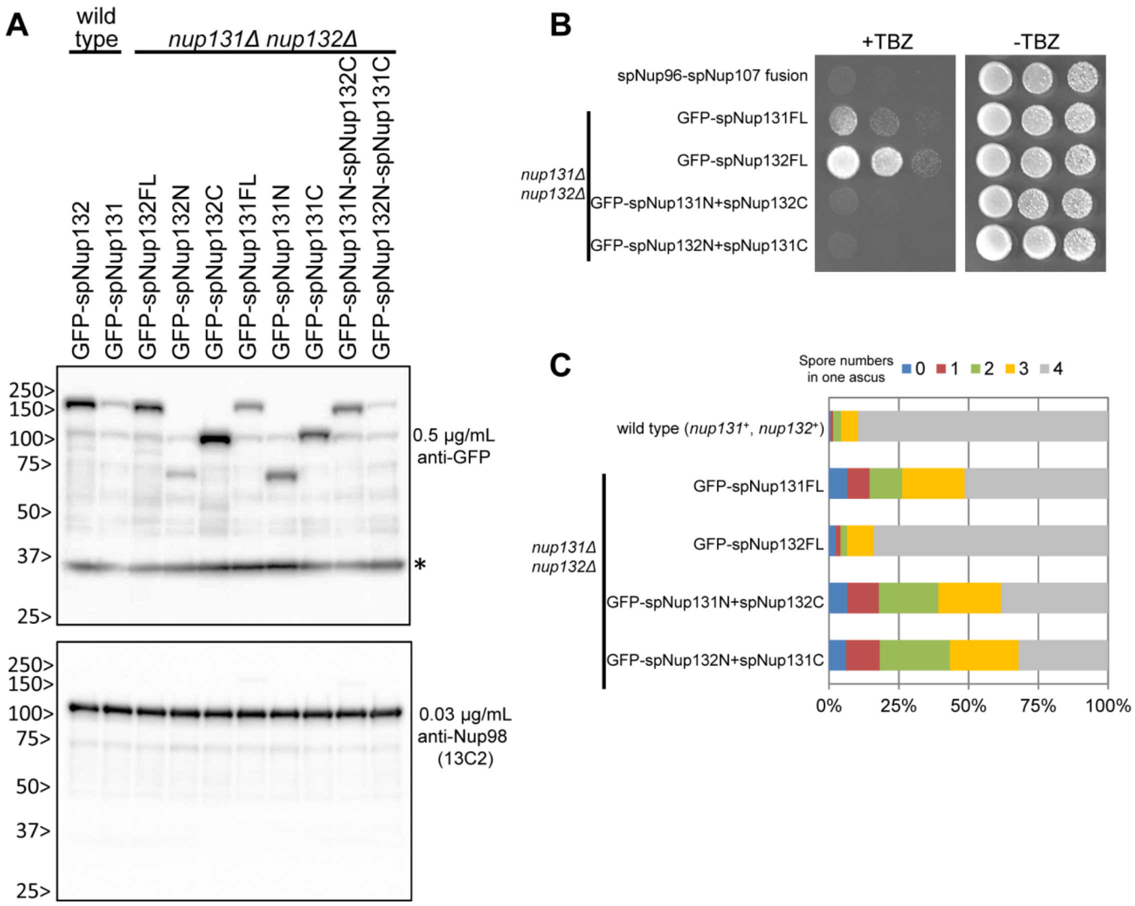


**S7 Fig. Characterization of the strains used in Fig. 6**

**(A)** Detection of GFP fused protein fragments by Western blot. *nup131*Δ *nup132*Δ cells expressing the indicated protein domains or chimeric proteins were analyzed by Western blot. Extracts prepared from wild type strains expressing GFP-spNup132 and GFP-spNup131 were applied to the left two lanes to examine the expression level of the endogenous proteins. Asterisk indicates a non-specific cross reaction of the anti-GFP antibody. spNup98 was detected with the anti-Nup98 antibody (13C2) as a loading control. **(B)** TBZ sensitivity of *nup131*Δ *nup132*Δ cells expressing the indicated protein domains or chimeric proteins. Five-fold serial dilutions of cells indicated were spotted on YES medium in the presence (+TBZ) or absence (-TBZ) of TBZ (used at 20μg/mL in this experiment). The plates were observed after 3-5 days incubation. **(C)** Spore formation of *nup131*Δ *nup132*Δ cells expressing the indicated proteins. The spore number in each ascus is indicated. More than 200 zygotes were counted for each strain.

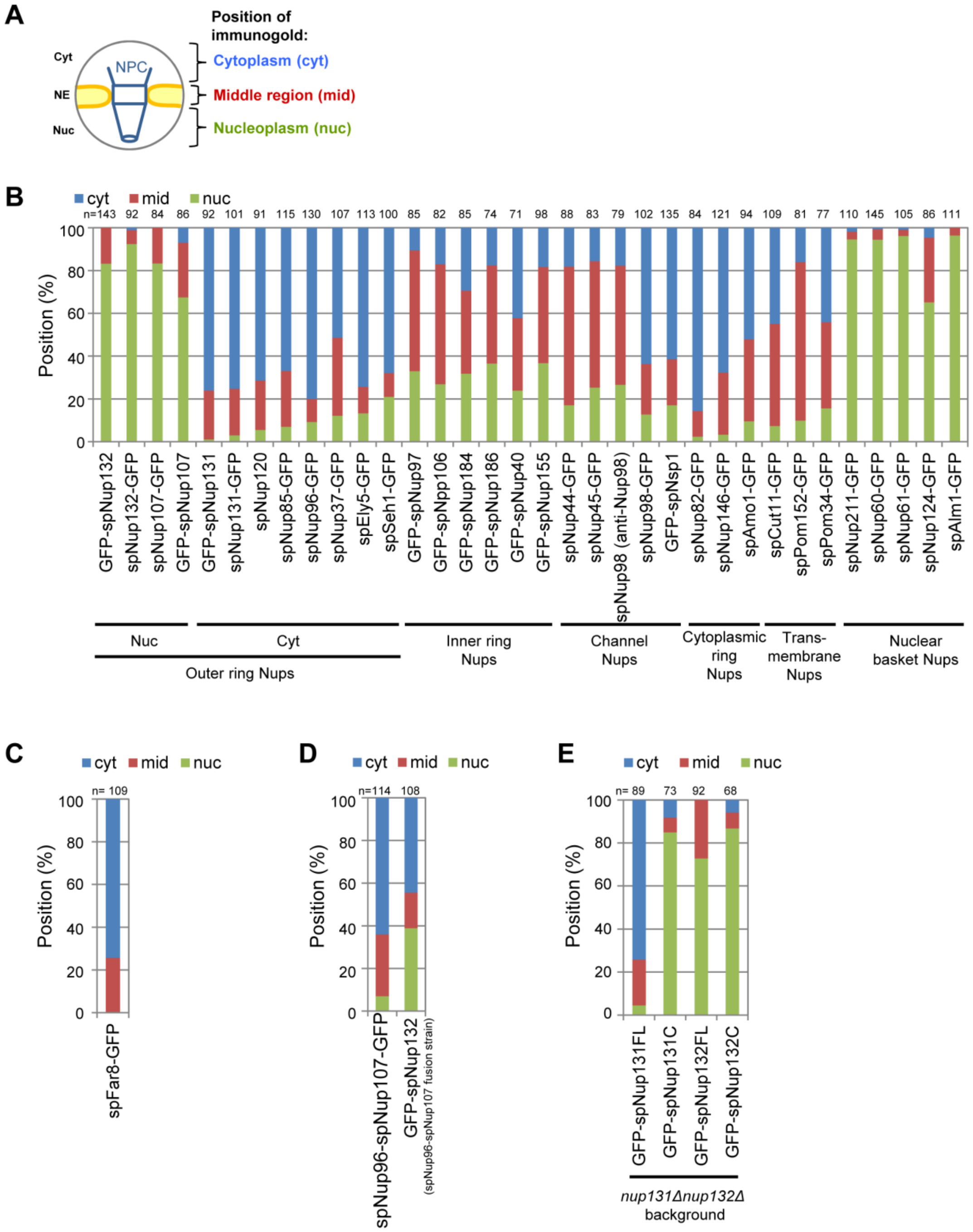


**S8 Fig. Distribution of Nups within the NPC**

**(A)** The schematic drawing indicates the cytoplasmic, middle and nuclear regions within the NPC. **(B)** Distribution of GFP fused proteins indicated at the bottom. The number of immuno-gold particles in the IEM image data (Figs 1B-C, 3A, 4A, and 7A-E) was counted for each of the NPC regions. The colored bar graph indicates percentages of each region. The total numbers of gold particles counted are indicated at the top of each column. **(C)** IEM image data of Fig 2B were analyzed. **(D)** IEM image data of Fig 5B-C were analyzed. **(E)** IEM image data of Fig 6C were analyzed.

### S1 Table. Nucleoporins in *S. pombe*, *S. cerevisiae*, and *H. sapiens**

| Subcomplex | *S. pombe* | *S. cerevisiae* | *H. sapiens* |
|---|---|---|---|
| Transmembrane Nups | spCut11 | scNdc1 | hsNdc1 |
| | spPom34/Mug31 | scPom34 | — |
| | spPom152 | scPom152 | — |
| | spTts1 | scPom33 | hsTMEM33 |
| | | scPer33 | |
| | — | — | hsGp210/Nup210 |
| | — | — | hsPom121 |
| Outer ring Nups | spEly5 | — | hsELYS |
| | spNup37 | — | hsNup37 |
| | spNup85 | scNup85 | hsNup85 |
| | spNup107 | scNup84 | hsNup107 |
| | spNup120 | scNup120 | hsNup160 |
| | spNup131 | scNup133 | hsNup133 |
| | spNup132 | | |
| | spNup189c | scNup145c | hsNup96 |
| | spSeh1 | scSeh1 | hsSeh1 |
| | —** | scSec13 | hsSec13 |
| | — | — | hsNup43 |
| Inner ring Nups | spNup97/Mug87 | scNic96 | hsNup93 |
| | spNpp106 | | |
| | spNup184 | scNup188 | hsNup188 |
| | spNup186 | scNup192 | hsNup205 |
| | spNup155 | scNup157 | hsNup155 |
| | | scNup170 | |
| | spNup40 | scNup53 | hsNup35 (MP-44) |
| | | scNup59 | |
| Channel Nups | spNsp1 | scNsp1 | hsNup62 |
| | spNup44 | scNup57 | hsNup54 |
| | spNup45 | scNup49 | hsNup58 |
| | spNup189n | scNup100 | hsNup98 |
| | | scNup116 | |
| | | scNup145n | |
| Cytoplasmic Nups | spNup82 | scNup82 | hsNup88 |
| | spNup146 | scNup159 | hsNup214 |
| | spAmo1 | scNup42/Rip1 | hsNlp1/hCG1/NUPL2 |
| | — | — | hsALADIN |
| | — | — | hsNup358 |
| Nuclear basket Nups | spNup60 | scNup60 | - |
| | spNup61 | scNup2 | hsNup50 |
| | spNup124 | scNup1 | hsNup153 |
| | spAlm1 | — | — |
| | spNup211 | scMlp1 | hsTpr |
| | | scMlp2 | |

*This Table is modified from Asakawa *et al*. 2014.

**Sec13 does not show nuclear periphery localization in *S. pombe*, thus it is not included in the "*S. pombe*" column and is represented as (-) (Asakawa *et al*. 2014).

### S2 Table. *S. pombe* strains used in this study

| Figs | Description | Strain name | Genotypes | Reference |
|---|---|---|---|---|
| Fig.1, S3Fig | GFP-spNup131 | HA1133 | $h^{90}$ *ade6-216 ura4 leu1-32*<br>*lys1*$^{+}$*::Pnup131-GFP-nup131*$^{+}$ *nup131::ura4*$^{+}$ | Asakawa<br>et al. 2014 |
| Fig.1,<br>S2D Fig,<br>S3 Fig,<br>Fig.5C,<br>S8B Fig | GFP-spNup132 | HA1131 | $h^{90}$ *ade6-216 ura4 leu1-32*<br>*lys1*$^{+}$*::Pnup132-GFP-nup132*$^{+}$ *nup132::ura4*$^{+}$ | Asakawa<br>et al. 2014 |
| Fig.1,<br>S2B,C Fig,<br>S8B Fig. | GFP-spNup131<br>spMis6-GFP | HA1374-9C | $h^{90}$ *ade6-216 ura4 leu1-32*<br>*lys1*$^{+}$*::Pnup131-GFP-nup131*$^{+}$ *nup131::ura4*$^{+}$<br>*mis6-GFP-LEU2* | This study |
| Fig.1,<br>S8B Fig. | spNup131-GFP | H03/C12 | $h^{90}$ *ade6-216 ura4 leu1-32 lys1-131*<br>*nup131-GFP-HA-kan*$^{r}$ | Asakawa<br>et al. 2014 |
| Fig.1,<br>S8B Fig. | spNup132-GFP | H02/C10 | $h^{90}$ *ade6-216 ura4 leu1-32 lys1-131*<br>*nup132-GFP-HA-kan*$^{r}$ | Asakawa<br>et al. 2014 |
| Fig.1 | GFP-spNup131<br>mCherry-spNup132 | HA1619 | $h^{90}$ *ade6-216 ura4-D18 leu1-32 nup131::ura4+*<br>*nup132::ura4*$^{+}$ *lys1*$^{+}$*::GFP-nup131*$^{+}$<br>*aur1*$^{r}$*::mCherry-nup132*$^{+}$ | This study |
| Fig.1 | mCherry-spNup131<br>GFP-spNup132 | HA1618 | $h^{90}$ *ade6-216 ura4-D18 leu1-32 nup131::ura4*$^{+}$<br>*nup132::GFP-nup132*$^{+}$ *lys1*$^{+}$*::mCherry-nup131*$^{+}$ | This study |
| S2A Fig. | spMis6-GFP | CRLc27 | $h^{-}$ *leu1-32 lys1-131 ade6-216 mis6-GFP::LEU2* | This study |
| Fig.2A,C | wild type<br>spFar8-GFP<br>spCut11-mCherry | HA1932 | $h^{-}$ *cut11-mCherry-hph far8-GFP-nat* | This study |
| Fig.2A,C | *nup131Δ*<br>spFar8-GFP<br>spCut11-mCherry | HA1940 | $h^{-}$ *nup131::kan*$^{r}$ *cut11-mCherry-hph far8-GFP-nat* | This study |
| Fig.2A,C | *nup132Δ*<br>spFar8-GFP<br>spCut11-mCherry | HA1933 | $h^{-}$ *nup132::kan*$^{r}$ *cut11-mCherry-hph far8-GFP-nat* | This study |
| Fig.2B,<br>S8C Fig. | spFar8-GFP | HA1824-9B | $h^{-}$ *lys1-131 far8-GFP-nat* | This study |
| Fig.2D,E | Vector | HA2030 | $h^{90}$ *ade6-216 ura4-D18 leu1-32*<br>*lys1*$^{+}$*::pCST3(empty vector) nup131::ura4*$^{+}$<br>*cut11-mCherry-hph far8-GFP-kan*$^{r}$ | This study |
| Fig.2D,E | spNup131op | HA2031 | $h^{90}$ *ade6-216 ura4-D18 leu1-32*<br>*lys1*$^{+}$*::Pnmt1-nup131*$^{+}$ *nup131::ura4*$^{+}$<br>*cut11-mCherry-hph far8-GFP-kan*$^{r}$ | This study |
| Fig.2D,E | spNup132op | HA2032 | *h90 ade6-216 ura4-D18 leu1-32*<br>*lys1+::Pnmt1-nup132+ nup131::ura4+*<br>*cut11-mCherry-hph far8-GFP-kanr* | This study |
| S4 Fig. | wild type<br>spFar11-GFP<br>spCut11-mCherry | HA1936 | $h^{-}$ *cut11-mCherry-hph far11-GFP-nat* | This study |
| S4 Fig. | *nup131Δ*<br>spFar11-GFP<br>spCut11-mCherry | HA1942 | $h^{-}$ *nup131::kan*$^{r}$ *cut11-mCherry-hph*<br>*far11-GFP-nat* | This study |
| S4 Fig. | *nup132Δ*<br>spFar11-GFP<br>spCut11-mCherry | HA1937 | $h^{-}$ *nup132::kan*$^{r}$ *cut11-mCherry-hph*<br>*far11-GFP-nat* | This study |
| Fig.3A,<br>S8B Fig. | spNup211-GFP | YN023-2-2<br>A | $h^{90}$ *leu1-32 lys1-131 ura4-D18 ade6-216*<br>*nup211-GFP::ura4*$^{+}$ | Asakawa<br>et al. 2014 |

| | | | | |
|---|---|---|---|---|
| Fig.3B,C | wild type spNup211-GFP spCut11-mCherry | HA1833-1C | *h*$^-$ *ura4 lys1 cut11-mCherry-hph nup211-GFP-ura4*$^+$ | This study |
| Fig.3B,C | *nup131*Δ spNup211-GFP spCut11-mCherry | HA1832-1A | *h*$^-$ *ura4 lys1 nup131::kanr cut11-mCherry-hph nup211-GFP-ura4*$^+$ | This study |
| Fig.3B,C | *nup132*Δ spNup211-GFP spCut11-mCherry | HA1833-1B | *h*$^-$ *ura4 lys1 nup132::nat cut11-mCherry-hph nup211-GFP-ura4*$^+$ | This study |
| Fig. 3C | *nup132*Δ +vector | HA2040 | *h*$^-$ *ura4 lys1*$^+$*::pCST3 nup132::nat cut11-mCherry-hph nup211-GFP-ura4*$^+$ | This study |
| Fig. 3C | *nup132*Δ +*nup132*$^+$ | HA2042 | *h*$^-$ *ura4 lys1*$^+$*::Pnmt1-nup132*$^+$*(cDNA) nup132::nat cut11-mCherry-hph nup211-GFP-ura4*$^+$ | This study |
| Fig.4, S5,S8B Figs. | spNup120-GFP spMis6-GFP | HA1628 | *h*$^{90}$ *ade6-216 ura4-D18 leu1-32 lys1-131 nup120-GFP-HA-kanr mis6-GFP-hph* | This study |
| Fig.4, S5,S8B Figs. | spNup85-GFP spMis6-GFP | HA1629 | *h*$^{90}$ *ade6-216 ura4-D18 leu1-32 lys1-131 nup85-GFP-HA-kanr mis6-GFP-hph* | This study |
| Fig.4, S5,S8B Figs. | spNup96-GFP spMis6-GFP | HA1626 | *h*$^{90}$ *ade6-216 ura4-D18 leu1-32 lys1-131 nup189c-GFP-HA-kan*$^r$ *mis6-GFP-hph* | This study |
| Fig.4, S5,S8B Figs. | spEly5-GFP spMis6-GFP | HA1633 | *h*$^{90}$ *ade6-216 ura4-D18 leu1-32 lys1-131 ely5-GFP-HA-kanr mis6-GFP-hph* | This study |
| Fig.4, S5,S8B Figs. | spNu37-GFP spMis6-GFP | HA1627 | *h*$^{90}$ *ade6-216 ura4-D18 leu1-32 lys1-131 nup37-GFP-kan*$^r$ *mis6-GFP-hph* | This study |
| Fig.4, S5,S8B Figs. | spSeh1-GFP spMis6-GFP | HA1636 | *h*$^{90}$ *ade6-216 ura4-D18 leu1-32 lys1-131 seh1-GFP-HA-kan*$^r$ *mis6-GFP-hph* | This study |
| Fig.4, S5,S8B Figs. | spNup107-GFP | H16/D11 | *h*$^{90}$ *ade6-216 ura4-D18 leu1-32 lys1-131 nup107-GFP-HA-kan*$^r$ | Asakawa et al. 2014 |
| Fig.4, S8B Fig. | GFP-spNup107 | HA1657 | *h*$^{90}$ *ade6-216 ura4-D18 leu1-32 lys1-131 GFP-nup107-kan*$^r$ | This study |
| Fig.4 | spNup85-GFP mCherry-spNup131 | HA1965 | *h*$^{90}$ *ade6-216 ura4-D18 leu1-32 nup131::ura4*$^+$ *lys1*$^+$*::Pnup131-mCherry-nup131 nup85-GFP-kan*$^r$ | This study |
| Fig.4 | spNup85-GFP mCherry-spNup132 | HA1966 | *h*$^{90}$ *ade6-216 ura4-D18 leu1-32 lys1-131 nup132::ura4*$^+$ *aur1*$^r$*::Pnup132-mCherry-nup132 nup85-GFP-kan*$^r$ | This study |
| Fig.4 | spNup107-GFP mCherry-spNup131 | HA2020 | *h*$^{90}$ *ade6-216 ura4-D18 leu1-32 nup131::ura4*$^+$ *lys1*$^+$*::Pnup131-mCherry-nup131 nup107-GFP-hph* | This study |
| Fig.4 | spNup107-GFP mCherry-spNup132 | HA2021 | *h*$^{90}$ *ade6-216 ura4-D18 leu1-32 nup132::ura4*$^+$ *lys1*$^+$*::Pnup132-mCherry-nup132 nup96-GFP-hph* | This study |
| Fig.5A,D,G | wild type | AY160-14D | *h*$^{90}$ *ade6-216 ura4 leu1-32 lys1-131* | Hayashi et al. 2009 |
| Fig.5A | spNup96-spNup107 -GFP | HA1774-2B | *h*$^{90}$ *ade6-216 ura4 leu1-32 lys1-131 nup189::FL5-nup107full-GFP-kanr nup107::nat* | This study |
| Fig.5B, S5, S8D Figs. | spNup96-spNup107 -GFP spMis6-GFP | HA1905 | *h*$^{90}$ *ade6-216 ura4 leu1-32 lys1-131 nup189::FL5-nup107full-GFP-kanr nup107::nat mis6-GFP-hph* | This study |
| Fig.5C, S5, S8D Figs. | spNup96-spNup107 fusion GFP-spNup132 | HA1783-14 C | *h*$^{90}$ *ade6-216 ura4 leu1 lys1*$^+$*::GFP-nup132 nup132::ura4*$^+$ *nup189*$^+$*::FL5-nup107full-hph nup107::nat* | This study |
| Fig.5D,G | spNup96-spNup107 fusion | HA1776-6A | *h*$^{90}$ *ade6-216 ura4 leu1 lys1 nup189*$^+$*::FL5-nup107full-hph nup107::nat* | This study |
| Fig.5E,F, | wild type | HA1181 | *h*$^{90}$ *ade6-210 leu1-32 lys1-131 ura4-D18* | This study |

| | | | | |
|---|---|---|---|---|
| S6 Fig. | mCherry-spAtb2 | | aur1$^{r}$-nda3pro-mCherry-atb2$^{+}$ | |
| Fig.5E,F | spNup96-spNup107 fusion mCherry-spAtb2 | HJY896 | $h^{90}$ ade6-216 leu1-32 lys1-131 ura4-D18 nup189$^{+}$::FL5-nup107full-hph nup107::nat aur1$^{r}$::Pnda3-GFP-atb2$^{+}$ | This study |
| S6 Fig. | *nup131Δ* | HA2107 | $h^{90}$ ade6-210 ura4-D18 leu1-32 lys1-131 nup131::ura4$^{+}$ aur1$^{r}$::Pnda3-mCherry-atb2$^{+}$ | This study |
| Fig.6B, S7 Fig. | spNup132FL | HJY760 | $h^{90}$ ade6-216 ura4-D18 leu1-32 lys1-131 nup131::ura4$^{+}$ nup132::ura4$^{+}$ cut11-mCherry-hph lys1$^{+}$::Pnup132-GFPs65t-nup132FL-Tnmt1 | This study |
| Fig.6B, S7 Fig. | spNup131FL | HJY773 | $h^{90}$ ade6-216 ura4-D18 leu1-32 lys1-131 nup131::ura4$^{+}$ nup132::ura4$^{+}$ cut11-mCherry-hph lys1$^{+}$-Pnup132-GFPs65t-nup131FL | This study |
| Fig.6B, S7 Fig. | spNup132C | HJY769 | $h^{90}$ ade6-216 ura4-D18 leu1-32 lys1-131 nup131::ura4$^{+}$ nup132::ura4$^{+}$ cut11-mCherry-hph lys1$^{+}$-Pnup132-GFPs65t-nup132C | This study |
| Fig.6B, S7 Fig. | spNup131C | HJY756 | $h^{90}$ ade6-216 ura4-D18 leu1-32 lys1-131 nup131::ura4$^{+}$ nup132::ura4$^{+}$ cut11-mCherry-hph lys1$^{+}$-Pnup132-GFPs65t-nup131C | This study |
| Fig.6B, S7 Fig. | spNup132N | HJY855 | $h^{90}$ ade6-216 ura4-D18 leu1-32 lys1-131 nup131::ura4$^{+}$ nup132::ura4$^{+}$ cut11-mCherry-hph lys1$^{+}$::Pnup132-GFPs65t-nup132N-Tnmt1 | This study |
| Fig.6B, S7 Fig. | spNup131N | HJY856 | $h^{90}$ ade6-216 ura4-D18 leu1-32 lys1-131 nup131::ura4$^{+}$ nup132::ura4$^{+}$ cut11-mCherry-hph lys1$^{+}$::Pnup132-GFPs65t-nup131N-Tnmt1 | This study |
| Fig.6B, S7 Fig. | spNup131N+spNup 132C | HJY763 | $h^{90}$ ade6-216 ura4-D18 leu1-32 lys1-131 nup131::ura4$^{+}$ nup132::ura4$^{+}$ cut11-mCherry-hph lys1$^{+}$-Pnup132-GFPs65t-nup131N-nup132C | This study |
| Fig.6B, S7 Fig. | spNup132N+spNup 131C | HJY771 | $h^{90}$ ade6-216 ura4-D18 leu1-32 lys1-131 nup131::ura4$^{+}$ nup132::ura4$^{+}$ cut11-mCherry-hph lys1$^{+}$-Pnup132-GFPs65t-nup132N-nup131C | This study |
| Fig.6C, S8E Fig. | GFP-spNup132FL | HJY728 | $h^{90}$ ade6-216 ura4-D18 leu1-32 lys1-131 nup131::ura4$^{+}$ nup132::ura4$^{+}$ lys1$^{+}$::Pnup132-GFPs65t-nup132FL-Tnmt1 | This study |
| Fig.6C, S8E Fig. | GFP-spNup132C | HJY782 | $h^{90}$ ade6-216 ura4-D18 leu1-32 lys1-131 nup131::ura4$^{+}$ nup132::ura4$^{+}$ lys1$^{+}$::Pnup132-GFPs65t-nup132L+C-Tnmt1 | This study |
| Fig.6C, S8E Fig. | GFP-spNup131FL | HJY810 | $h^{90}$ ade6-216 ura4-D18 leu1-32 lys1-131 nup131::ura4$^{+}$ nup132::ura4$^{+}$ lys1$^{+}$-Pnup132-GFPs65t-nup131FL | This study |
| Fig.6C, S8E Fig. | GFP-spNup131C | HJY743 | $h^{90}$ ade6-216 ura4-D18 leu1-32 lys1-131 nup131::ura4$^{+}$ nup132::ura4$^{+}$ lys1$^{+}$::Pnup132-GFPs65t-nup131L+C-Tnmt1 | This study |
| S7 Fig. | GFP-spNup132 | HA1523 | $h^{90}$ ade6-216 leu1-32 ura4-D18 lys1$^{+}$::Pnup132-GFP::nup132$^{+}$ nup132::ura4$^{+}$ cut11-mCherry-hph | This study |
| S7 Fig. | GFP-spNup131 | HA1524 | $h^{90}$ ade6-216 leu1-32 ura4-D18 lys1$^{+}$::Pnup131-GFP::nup131$^{+}$ nup131::ura4$^{+}$ cut11-mCherry-hph | This study |
| S7 Fig. | wild type | HA2128 | $h^{90}$ ade6-216 ura4-D18 leu1-32 lys1$^{+}$::pCSU3 cut11-mCherry-hph | This study |
| Fig.7, S5, S8 Figs | GFP-spNup97 | HA1330-1C | $h^{90}$ ade6-216 ura4-D18 leu1-32 nup97::LEU2 lys1$^{+}$::GFP-nup97 | Asakawa et al. 2014 |
| Fig.7, S5, S8 Figs | GFP-spNpp106 | HA1658 | $h^{90}$ ade6-216 ura4-D18 leu1-32 lys1-131 GFP-npp106-kanr | This study |

| | | | | |
|---|---|---|---|---|
| Fig.7, S5, S8 Figs | GFP-spNup184 | HA1656 | $h^{90}$ *ade6-216 ura4-D18 leu1-32 lys1-131 GFP-nup184-kan*$^{r}$ | This study |
| Fig.7, S5, S8 Figs | GFP-spNup186 | HA1659 | $h^{90}$ *ade6-216 ura4-D18 leu1-32 lys1-131 GFP-nup186-kan*$^{r}$ | This study |
| Fig.7, S5, S8 Figs | GFP-spNup40 | HA1134 | $h^{90}$ *ade6-216 ura4-D18 leu1-32 lys1*$^{+}$*::Pnup40-GFP-nup40*$^{+}$ *nup40::ura4*$^{+}$ | Asakawa et al. 2014 |
| Fig.7, S5, S8 Figs | spNup155-GFP | AHP001 | $h^{90}$ *ade6-216 ura4-D18 leu1-32 lys1-131 nup155-GFP-kan*$^{r}$ | Asakawa et al. 2014 |
| Fig.7, S5, S8 Figs | spNup44-GFP | H04/D06 | $h^{90}$ *ade6-216 ura4-D18 leu1-32 lys1-131 nup44-GFP-HA-kan*$^{r}$ | Asakawa et al. 2014 |
| Fig.7, S5, S8 Figs | spNup45-GFP | HA1622 | $h^{90}$ *ade6-216 ura4 leu1-32 lys1-131 nup45-GFP-kan*$^{r}$ | Asakawa et al. 2014 |
| Fig.7, S5, S8 Figs | spNup98-GFP | HA800-7A | $h^{90}$ *ade6-216 ura4-D18 leu1-32 lys1-131 nup189N-GFP-kan*$^{r}$ | Asakawa et al. 2014 |
| Fig.7, S5, S8 Figs | spNup98 (anti-Nup98) | AY160-14D | $h^{90}$ *ade6-216 ura4 leu1-32 lys1-131* | Hayashi et al. 2009 |
| Fig.7, S5, S8 Figs | GFP-spNsp1 | AHP018 | $h^{90}$ *GFP-nsp1-kan*$^{r}$ *ade6-216 ura4-D18 leu1-32 lys1-131* | Asakawa et al. 2014 |
| Fig.7, S5, S8 Figs | spNup82-GFP | H04/H10 | $h^{90}$ *nup82-GFP-HA-kan*$^{r}$ *ade6-216 ura4-D18 leu1-32 lys1-131* | Asakawa et al. 2014 |
| Fig.7, S5, S8 Figs | spNup146-GFP | H03/D05 | $h^{90}$ *ade6-216 ura4-D18 leu1-32 lys1-131 nup146-GFP-HA-kan*$^{r}$ | Asakawa et al. 2014 |
| Fig.7, S5, S8 Figs | spAmo1-GFP | HA1062-4C | $h^{90}$ *ade6-216 ura4-D18 leu1-32 lys1-131 amo1-GFP-kan*$^{r}$ | Asakawa et al. 2014 |
| Fig.7, S5, S8 Figs | spCut11-GFP | H14/C04 | $h^{90}$ *ade6-216 ura4-D18 leu1-32 lys1-131 cut11-GFP-HA-kan*$^{r}$ | Asakawa et al. 2014 |
| Fig.7, S5, S8 Figs | spPom152-GFP | H04/D07 | $h^{90}$ *pom152-GFP-HA-kan*$^{r}$ *ade6-216 ura4-D18 leu1-32 lys1-131* | Asakawa et al. 2014 |
| Fig.7, S5, S8 Figs | spPom34-GFP | AHP016 | $h^{90}$ *ade6-216 ura4-D18 leu1-32 lys1-131 pom34-GFP-kan*$^{r}$ | Asakawa et al. 2014 |
| Fig.7, S5, S8 Figs | spNup60-GFP | AHP011 | $h^{90}$ *nup60-GFP-kan*$^{r}$ *ade6-216 ura4-D18 leu1-32 lys1-131* | Asakawa et al. 2014 |
| Fig.7, S5, S8 Figs | GFP-spNup61 | HA1132 | $h^{90}$ *ade6-216 ura4-D18 leu1-32 lys1*$^{+}$*::Pnup61-GFP::nup61*$^{+}$ *nup61::ura4*$^{+}$ | Asakawa et al. 2014 |
| Fig.7, S5, S8 Figs | GFP-spNup124 | HA1135 | $h^{90}$ *ade6-216 ura4-D18 leu1-32 lys1*$^{+}$*::Pnup124-GFP::nup124*$^{+}$ *nup124::ura4*$^{+}$ | Asakawa et al. 2014 |

,

**S3 Table. Dilution ratios of primary and secondary antibodies used for IEM**

| Figure | Strain | Primary antibody dilution* | Secondary antibody dilution* |
|---|---|---|---|
| Fig 1 | GFP-spNup131<br>GFP-spNup131 spMis6-GFP<br>spNup131-GFP<br>spNup132-GFP | 1:400 | 1:400 |
| | GFP-spNup132 | 1:400 | 1:100 |
| Fig 2B | spFar8-GFP | 1:400 | 1:400 |
| Fig 3A | spNup211-GFP | 1:400 | 1:400 |
| Fig 4A | spNup120-GFP spMis6-GFP<br>spNup85-GFP spMis6-GFP<br>spNup37-GFP spMis6-GFP | 1:400 | 1:200 |
| | spNup96-GFP spMis6-GFP<br>spEly5-GFP spMis6-GFP<br>spSeh1-GFP spMis6-GFP | 1:1000 | 1:800 |
| | spNup107-GFP<br>GFP-spNup107 | 1:400 | 1:400 |
| Fig 5B | spNup96-spNup107-GFP spMis6-GFP | 1:400 | 1:400 |
| Fig 5C | GFP-spNup132 | 1:400 | 1:400 |
| Fig 6C | GFP-spNup131FL<br>GFP-spNup131C<br>GFP-spNup132FL | 1:400 | 1:400 |
| | GFP-spNup132C | 1:400 | 1:1000 |
| Fig 7A | GFP-spNup97 | 1:400 | 1:200 |
| | GFP-spNup184 | 1:400 | 1:800 |
| | GFP-spNpp106<br>GFPspNup186<br>GFP-spNup40<br>spNup155-GFP | 1:400 | 1:400 |
| Fig 7B | spNup44-GFP<br>spNup45-GFP<br>spNup98-GFP<br>GFP-spNsp1 | 1:400 | 1:400 |
| Fig 7C | spNup82-GFP<br>spNup146-GFP<br>spAmo1-GFP | 1:400 | 1:400 |
| Fig 7D | spCut11-GFP | 1:400 | 1:200 |
| | spPom152-GFP<br>spPom34-GFP | 1:400 | 1:400 |
| Fig 7E | spNup60-GFP<br>spNup61-GFP | 1:400 | 1:200 |
| | GFP-spNup124<br>spAlm1-GFP | 1:400 | 1:400 |

*A rabbit polyclonal anti-GFP antibody (600-401-215, Rockland Immunochemicals, Limerick, PA, USA) was used as a primary antibody; a goat anti-rabbit Alexa 594 FluoroNanogold Fab' fragment (7304, Nanoprobes Inc., Yaphank, NY, USA) was used as a secondary antibody.
